# Stability and function of a putative microtubule organizing center in the human parasite *Toxoplasma gondii*

**DOI:** 10.1101/099267

**Authors:** Jacqueline M. Leung, Yudou He, Fangliang Zhang, Yu-Chen Hwang, Eiji Nagayasu, Jun Liu, John M. Murray, Ke Hu

## Abstract

The organization of the microtubule cytoskeleton is dictated by microtubule nucleators or organizing centers. *Toxoplasma gondii,* an important human parasite, has an array of 22 regularly spaced cortical microtubules stemming from a hypothesized organizing center, the apical polar ring. Here, we examine the functions of the apical polar ring by characterizing two of its components, KinesinA and APR1, and discovered that its putative role in templating can be separated from its mechanical stability. Parasites that lack both KinesinA and APR1 *(ΔkinesinAΔapr1)* are capable of generating 22 cortical microtubules. However, the apical polar ring is fragmented in live *ΔkinesinAΔapr1* parasites, and is undetectable by electron microscopy after detergent extraction. Disintegration of the apical polar ring results in the detachment of groups of microtubules from the apical end of the parasite. These structural defects are linked to a diminished ability of the parasite to move and to invade host cells, as well as decreased secretion of effectors important for these processes. Together, the findings demonstrate the importance of the structural integrity of the apical polar ring and the microtubule array in the *Toxoplasma* lytic cycle, which is responsible for massive tissue destruction in acute toxoplasmosis.

## INTRODUCTION

The cytoskeleton provides the framework for supporting numerous vital functions of a cell, including maintaining cell shape, providing mechanical strength, and driving material transport. Therefore, how a complex cytoskeletal structure is built and maintained from generation to generation is of general interest in biology. *Toxoplasma gondii,* an important human protozoan parasite, has a highly ordered and extraordinarily stereotyped cytoskeleton (Russell and Burns, 1984; Nichols and Chiappino, 1987; Hu *et al.,* 2002; Hu, 2008). This makes it an excellent system to understand the biophysical properties and principles of cytoskeletal assembly, and how these two elements are linked to cellular physiology. Furthermore, *T. gondii* is a powerful model organism for studying conserved aspects of biology in related, but less experimentally accessible members of the phylum Apicomplexa, including the *Plasmodium* spp. - the causative agents of malaria, *Cryptosporidium* spp., which cause cryptosporidiosis, and *Eimeria* spp., infectious agents of poultry and cattle.

Like other apicomplexans, *T. gondii* is an obligate intracellular parasite: it must navigate tissues, invade a host cell, replicate within, and lyse out of the host cell in order to survive. The parasite depends on its robust, yet flexible membrane cortex and underlying cytoskeleton to move and progress through the lytic cycle. The cytoskeleton includes several tubulin-based structures: a truncated cone-shaped structure (the “conoid”) made of 14 novel tubulin polymers wound in a left-handed spiral (Hu *et al.,* 2002), a pair of intra-conoid microtubules, and cortical (subpellicular) microtubules that extend in a left-handed spiral down two-thirds the length of the parasite (Nichols and Chiappino, 1987; Morrissette *et al.,* 1997) (Figure 1). During parasite replication, the cortical microtubules of the daughter parasites form inside the mother concomitantly with other components of the daughter scaffold, including a set of flattened vesicles sutured together (the inner membrane complex, or IMC) and an associated protein meshwork formed of intermediate-filament like proteins (Porchet and Torpier, 1977; Morrissette *et al.,* 1997). This scaffold provides the framework for housing the replicated organelles (the nucleus, Golgi, ER, apicoplast, and mitochondrion) and secretory organelles that are synthesized *de novo (e.g.* rhoptries, micronemes, and dense granules). Treatment of parasites with microtubule depolymerizing drugs inhibits the formation of new cortical microtubules and results in deformed, non-viable parasites (Morrissette and Sibley, 2002b; Morrissette *et al.,* 2004), suggesting that they direct the formation of the daughter scaffold. When fully polymerized, the cortical microtubules in mature parasites are resistant to microtubule-depolymerizing compounds, detergent extraction and low temperature conditions (Nichols and Chiappino, 1987; Morrissette *et al.,* 1997; Liu *et al.,* 2016). This extraordinary stability is conferred by coating proteins that heavily decorate the cortical microtubules (Hu *et al.,* 2002; Tran *et al.,* 2012; Liu *et al.,* 2013; Liu *et al.,* 2016).

**Figure 1.**
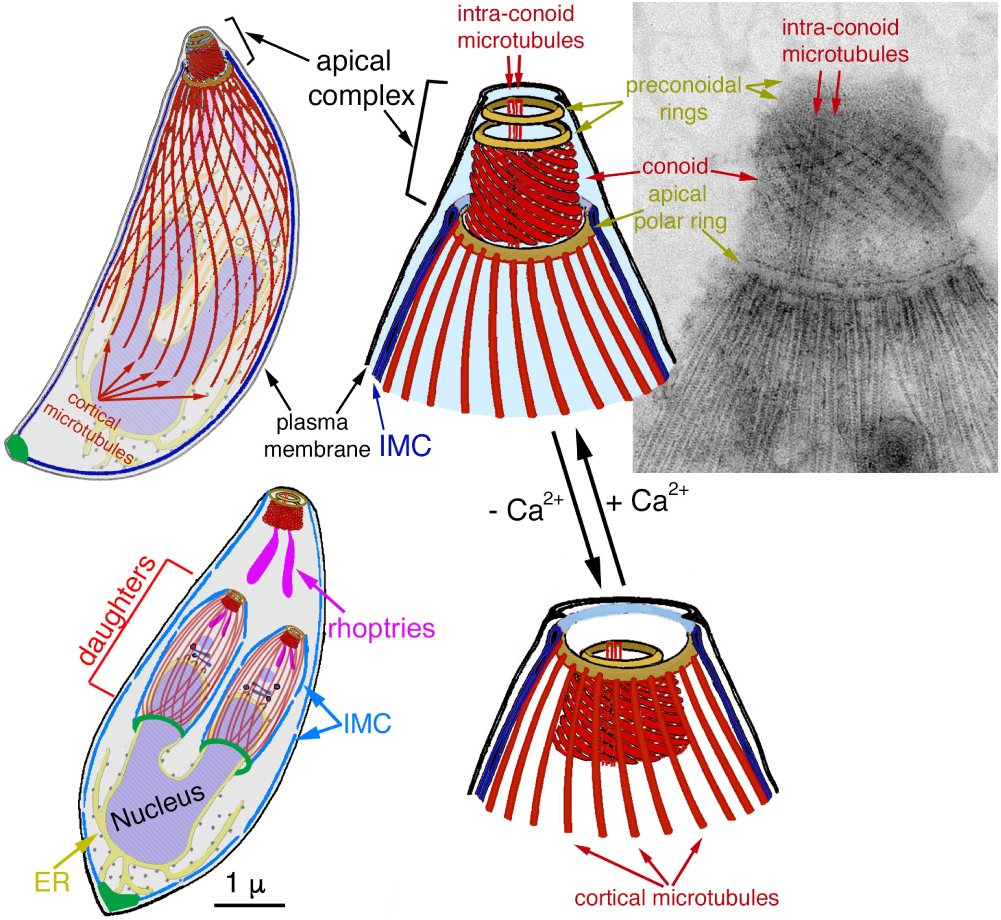
Cartoon and EM image depicting multiple tubulin-containing cytoskeletal structures (red) in *T. gondii,* including the 22 cortical microtubules, a pair of intra-conoid microtubules as well as the 14 fibers that make up the conoid. In its retracted state, the conoid forms a truncated cone that lies basal to the apical polar ring inside the parasite. Treatment of parasites with a calcium ionophore such as A23187 increases the concentration of calcium in the cytoplasm, inducing a change in the pitch of the conoid fibers, and protrusion of the conoid through the apical polar ring. Lower left, diagram showing the cortical microtubules forming during endodyogeny, a process where daughter parasites assemble inside the mother (mature) parasite. For clarity, only a subset of organelles are shown. The cortical microtubules of the mother parasite are not shown so those in the daughter parasites can be better visualized. IMC, inner membrane complex, ER, endoplasmic reticulum.

Ultrastructural studies showed that the cortical microtubules emanate from the apical polar ring (Figure 1) (Russell and Burns, 1984; Nichols and Chiappino, 1987; Bannister and Mitchell, 1995; Morrissette *et al.,* 1997). In *T. gondii* there are consistently 22 cortical microtubules, and their roots are evenly spaced between cogwheel-like projections at the ring circumference (Nichols and Chiappino, 1987). Its close apicomplexan relatives *Eimeria acervulina* and *Eimeria tenella* have 24 cortical microtubules during their sporozoite stage (Russell and Burns, 1984). In contrast, *Plasmodium* spp. asexual blood-stage parasites have a narrow band of 2-4 cortical microtubules that extends down one side of the parasite (Bannister and Mitchell, 1995; Morrissette and Sibley, 2002a). While there are variations in the number of cortical microtubules, the polymers in all of these cases are rooted at one end in the apical polar ring, which is thus thought to serve as their nucleating center. “Hook-decoration” (Heidemann and McIntosh, 1980) in extracted *Eimeria* parasites suggested that the apical polar ring might be attached to the minus (slow-growing) end of the microtubules (Russell and Burns, 1984) as is the case with microtubule organizing centers (MTOCs) in mammalian cells. Consistent with this view, growth of the cortical microtubules only occurs at the end distal to the apical polar ring (Hu *et al.,* 2002). However, it is not known what components of the apical polar ring dictate the exact number, precise spacing, orientation and polarity of these microtubules. It is also unclear how the apical polar ring provides a stable “root” for the cortical microtubules.

The apical polar ring is part of the cytoskeletal apical complex (Sheffield and Melton, 1968; Russell and Burns, 1984) that also includes the conoid, the intra-conoid microtubules and the preconoidal rings (Figure 1). The contents of specialized secretory organelles concentrated at the apical end of the parasite (the micronemes and rhoptries) are released through an opening at the apex of the conoid (Nichols *et al.,* 1983), to initiate formation of and mediate movement into a vacuole in which the parasite eventually resides. Discharge from these apical secretory organelles is important for parasite motility and invasion. One such example is micronemal protein 2 (MIC2), a protein secreted from the micronemes onto the parasite surface, serving as an adhesin that engages the parasite with its environment during motility and invasion into host cells (Huynh and Carruthers, 2006; Andenmatten *et al.,* 2012; Egarter *et al.,* 2014). Two components of the apical polar ring, RNG1 and RNG2, were identified previously (Tran *et al.,* 2010; Katris *et al.,* 2014). Although their structural roles are unknown, RNG2 was shown to function in constitutive and cGMP-stimulated secretion from the micronemes (Katris *et al.,* 2014).

Here, we show that the stability, but not the putative templating role, of the apical polar ring is dependent on two previously uncharacterized components, a putative kinesin (KinesinA), and a novel protein (<u>a</u>pical <u>p</u>olar <u>r</u>ing protein 1, APR1) without any known motifs. Upon the loss of KinesinA and APR1, the apical polar ring is fragmented in live parasites, resulting in the detachment of the cortical microtubules from the parasite apex, and consequently, irregular gaps in the array. These structural defects are connected with impaired parasite motility, secretion, and invasion, and a severe retardation of progression through the lytic cycle.

## RESULTS

### KinesinA and APR1 are differentially localized to and incorporated in the apical polar ring during daughter formation

We previously identified protein components of a fraction enriched for the cytoskeletal apical complex of *T. gondii* (Hu *et al.,* 2006). Localization of two of these proteins revealed new components of the apical polar ring, <u>a</u>pical <u>p</u>olar <u>r</u>ing protein 1 (APR1; TgGT1_315510, http://www.toxodb.org/toxo/, release 28) and a putative kinesin motor, KinesinA (TgGT1_267370). APR1 is conserved in closely related apicomplexan parasites, including the *Eimeria* spp., *Hammondia hammondi,* and *Neospora caninum.* but not in other apicomplexan lineages (Figure S1). The predicted motor domain (~aa37-430) at the N-terminus of KinesinA shows the highest degree of conservation in other apicomplexan parasites, including the *Plasmodium* spp., *Eimeria* spp., *H. hammondi,* and *N. caninum* (Figure S1).

To localize these proteins by fluorescence microscopy, we generated knock-in, endogenously tagged, and transgenic lines (Figures 2-5). Transgenic mCherryFP-tagged APR1 localized to a ring structure at the apical end of these parasites (Figure 2A). Immunoelectron microscopy of transgenic EGFP-tagged APR1 expressing parasites using an anti-GFP antibody generated a strong signal at the apical polar ring (Figure 2B) in good agreement with the light microscopy findings. Cytoplasmic aggregates are often seen in these transgenic lines (Figure 2A). This is likely an over-expression artifact, as they are not seen in *APR1-mCherryFP* knock-in parasites (Figure 4), generated using a previously described strategy (Heaslip *et al.,* 2010; Liu *et al.,* 2013; Liu *et al.,* 2016) in which the endogenous *apr1* gene was replaced with a LoxP-flanked APR1-mCherryFP expression cassette. Using live super-resolution structured illumination microscopy (SIM), KinesinA endogenously tagged at the C-terminus (KinesinA-mNeonGreenFP) was localized to a ring-shaped structure apical to APR1 (Figure 2A). Localization of KinesinA to the apical polar ring was further confirmed by treating the parasites with the calcium ionophore A23187 to induce conoid protrusion: since the KinesinA ring remained at the base of the conoid after protrusion, it could not be located at the preconoidal rings or the conoid, and thus it most likely is a component of the apical polar ring (Figure 2C). In contrast, another kinesin identified in an apical complex-depleted cytoskeletal fraction (Hu *et al.,* 2006), KinesinB (TgGT1_273560), was localized to the distal two-thirds of the cortical microtubules (Figure 2D), suggesting distinct roles for these two kinesins in *T. gondii.*

**Figure 2.**
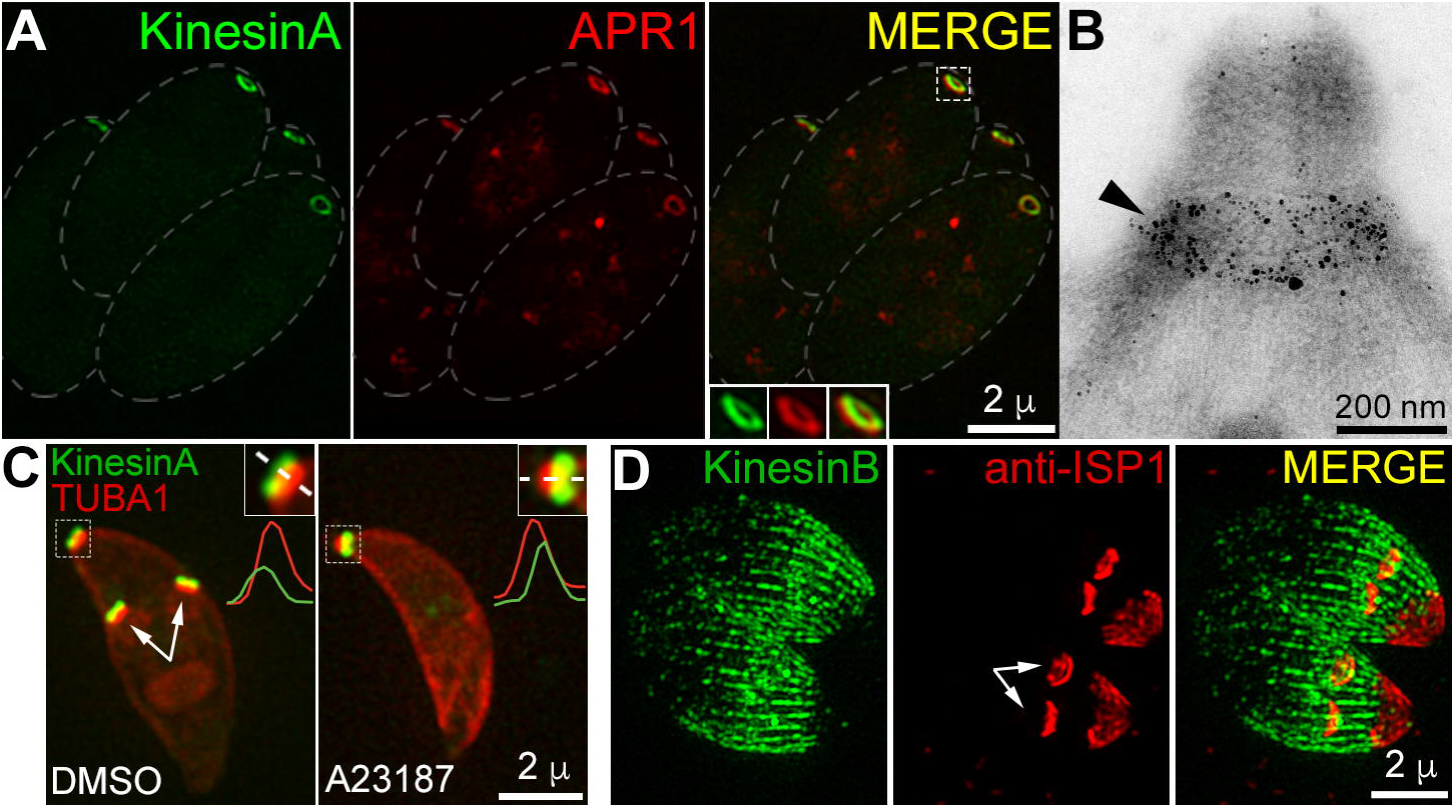
KinesinA and APR1 are localized to a ring structure at the apex of the parasite, with KinesinA located apical to APR1. **A.** 3D-structured illumination microscopy (SIM) projections of live intracellular parasites ectopically expressing ptubA1-APR1-mCherryFP (red) and KinesinA endogenously tagged with mNeonGreenFP (green). The ring that KinesinA-mNeonGreenFP forms is apical to that formed by APR1-mCherryFP. Insets are shown at 2X. **B.** Immunoelectron micrograph of parasites ectopically expressing EGFP-tagged APR1, extracted with sodium deoxycholate, labelled with anti-GFP antibody, 1.4 nm gold-secondary antibody, and silver enhanced. The gold/silver deposits are found in close proximity to the apical polar ring (black arrowhead). **C.** Wide-field epifluorescence images of live parasites showing that KinesinA (green) is localized apical to the conoid [labelled by transiently expressing mCherryFP-α1-tubulin (TUBA1, red) from a *T. gondii* tubulin promoter] when the conoid is retracted (DMSO, parasite treated with vehicle only). When the conoid is protruded in the presence of the calcium ionophore A23187, KinesinA is basal to the conoid. Quantification of the intensity profiles for both the KinesinA and TUBA1 signals along the dashed line is shown below the insets. Note that the parasite in the DMSO panel is replicating, and KinesinA is present in the apical complexes of the two developing daughters (arrows). Insets are at 2X. **D.** 3D-SIM projections of intracellular *KinesinB-mNeonGreenFP* endogenously tagged parasites, fixed and labelled with anti-ISP1 antibody. The apical boundary of KinesinB-mNeonGreenFP along the cortical microtubules coincides with the base of the apical cap marked by ISP1. Arrows indicate the ISP1 signal in daughters.

As a first step in investigating the functions of these new apical polar ring components, we determined the stages at which KinesinA and APR1 are present during daughter replication and assembly of the daughter cytoskeleton, based on incorporation of fluorescent tubulin subunits into the growing cortical microtubules (Hu *et al.,* 2002; Liu *et al.,* 2016). KinesinA-mNeonGreenFP was present in daughter parasites from the appearance of nascent daughters, through spindle pole formation, elongation of the daughter cytoskeleton, and emergence of fully formed daughters from the mother (Figure 3). In contrast, in *APR1-mCherryFP* knock-in parasites the mCherryFP signal was detectable in daughter parasites only during late stages of assembly, and daughter parasites as they emerged from the mother parasite (Figure 4). The spatial and temporal differences in the incorporation of these two components into the apical polar ring suggest that they may have non-overlapping roles in the parasite.

**Figure 3.**
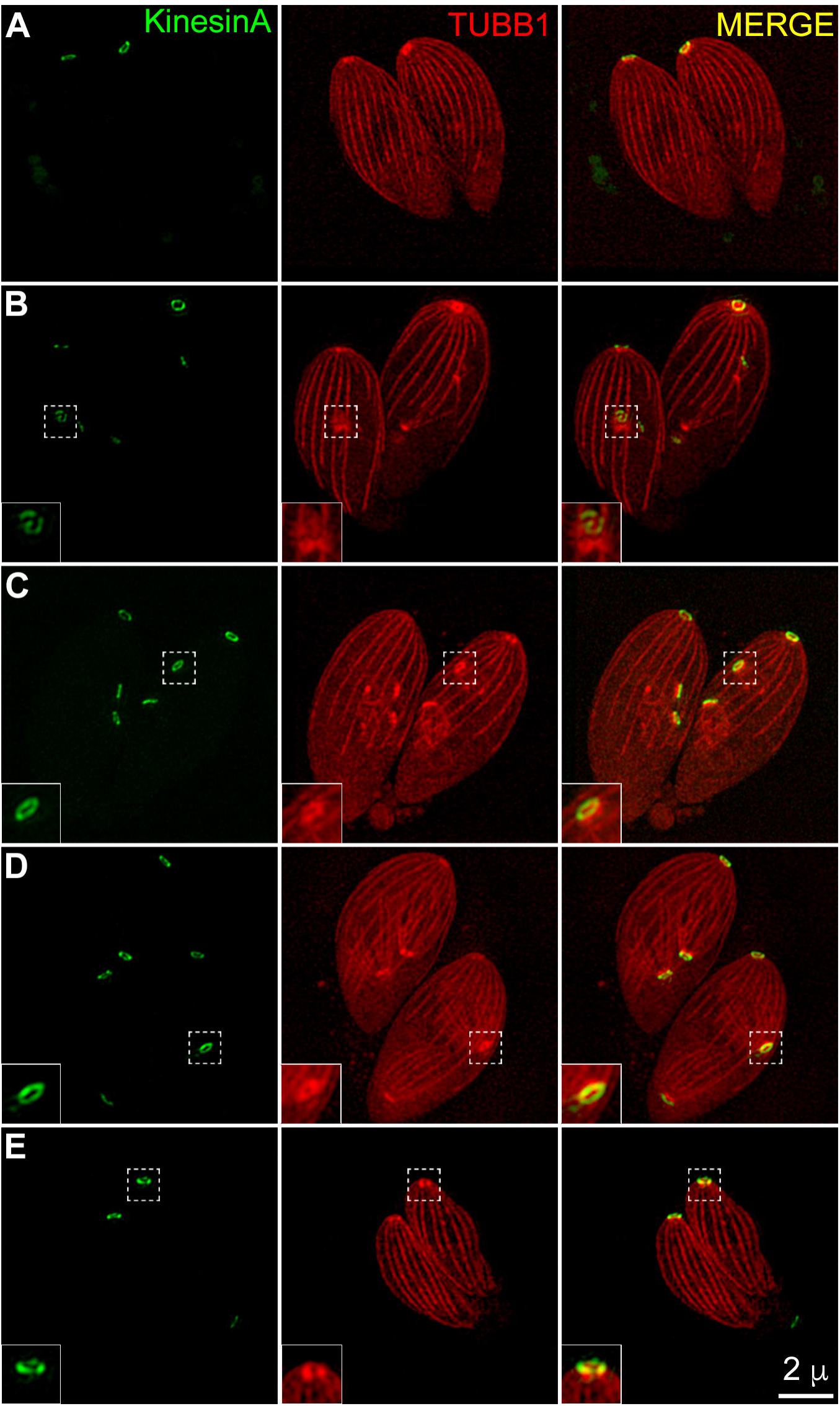
KinesinA is recruited to the apical polar ring during an early stage of daughter construction. Montage showing 3D-SIM projections of live, intracellular *KinesinA-mNeonGreenFP* (green) endogenously tagged parasites transiently expressing mAppleFP-β1-tubulin (TUBB1, red) from a *T. gondii* tubulin promoter. KinesinA-mNeonGreenFP is localized to the apical polar ring of mature, interphase parasites (A), and in daughters from the initiation of daughter formation (B) to the final stages as the fully assembled daughters emerge from the mother (E), of which only the residual body containing the mother apical polar ring is left. Insets show the apical region of one of the daughter parasites, enlarged at 2X.

**Figure 4.**
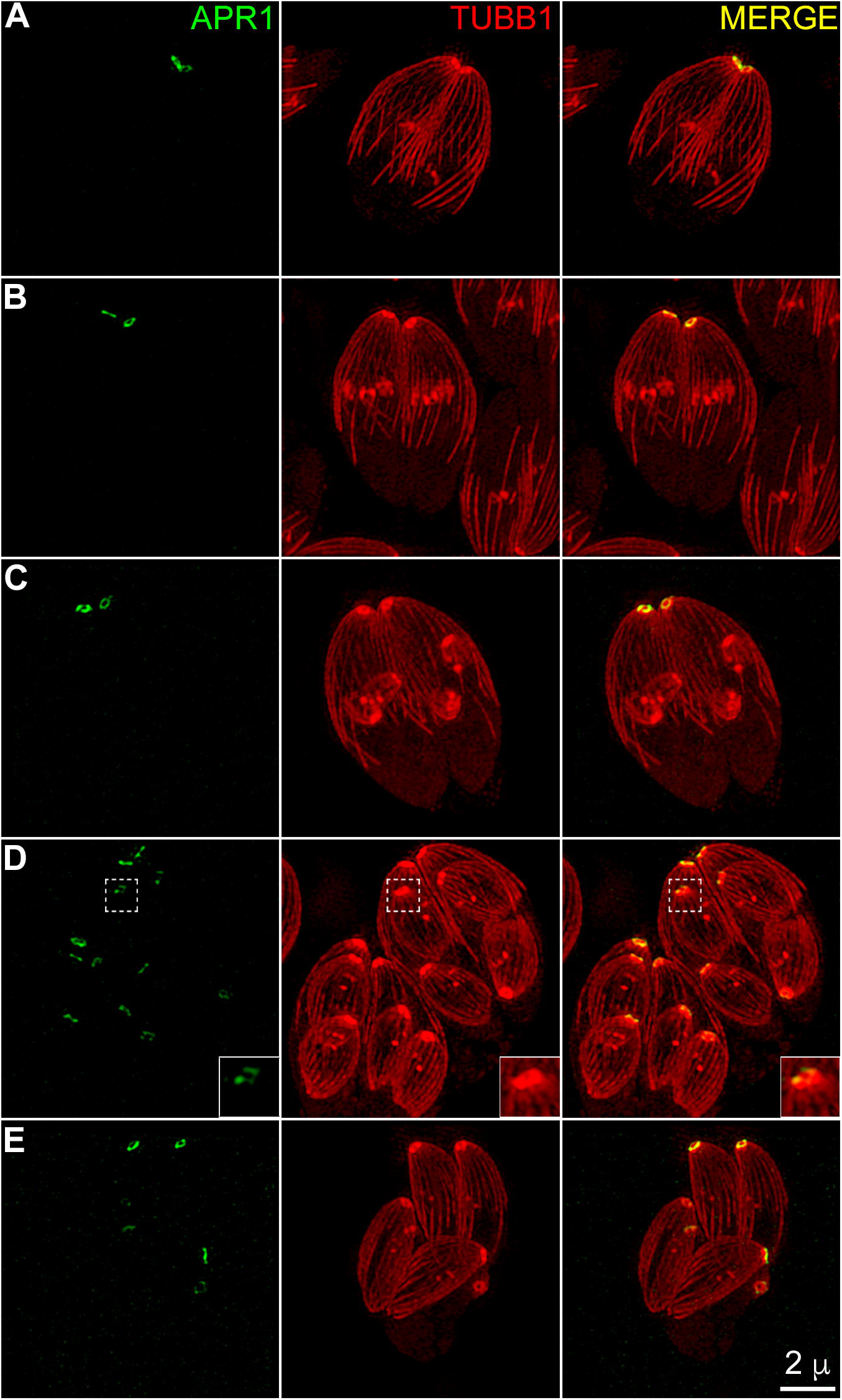
APR1 is recruited to the apical polar ring at a late stage of daughter construction. Montage showing 3D-SIM projections of live, intracellular *APR1-mCherryFP* (pseudocoloured green) knock-in parasites ectopically expressing EGFP-β1-tubulin (TUBB1, pseudocoloured red) from a *T. gondii* tubulin promoter (Hu, 2003; Wu *et al.,* 2016). APR1-mCherryFP is localized to the apical polar ring of mature, interphase parasites (A), and detected in daughters only during late stages of assembly and emergence from the mother (D & E). Insets show the apical region of one of the daughter parasites, enlarged at 2X.

### KinesinA and APR1 act synergistically in the parasite lytic cycle

To investigate the importance of these new apical polar ring components, we generated knockout mutants for *kinesinA* and *apr1* (Figure 5A and 5B). Due to the large size of the predicted *kinesinA* genomic locus (11,845 bp), knockout *(ΔkinesinA)* parasites were generated using the CRISPR/Cas9 system (Shen *et al.,* 2014; Sidik *et al.,* 2014). The endogenously tagged parasites described above were co-transfected with a *kinesinA*-targeting single guide RNA (sgRNA)-coding and Cas9 nuclease-coding plasmid, and a repair plasmid containing the 5’ and 3’ genomic regions of *kinesinA* but lacking its coding sequence in between (Figure 5A). Parasites were screened for the loss of the mNeonGreenFP fluorescence signal. The parental, *KinesinA-mNeonGreenFP* and *ΔkinesinA* parasites were confirmed by Southern blotting analysis. The genomic locus of *kinesinA* was also sequenced in the *ΔkinesinA* parasites to confirm the loss of the entire *kinesinA* gene (data not shown). The *ΔkinesinA* parasites had a modest growth defect compared to the wild-type or *KinesinA-mNeonGreenFP* parasites when assessed for plaquing efficiency (Figure 5C).

**Figure 5.**
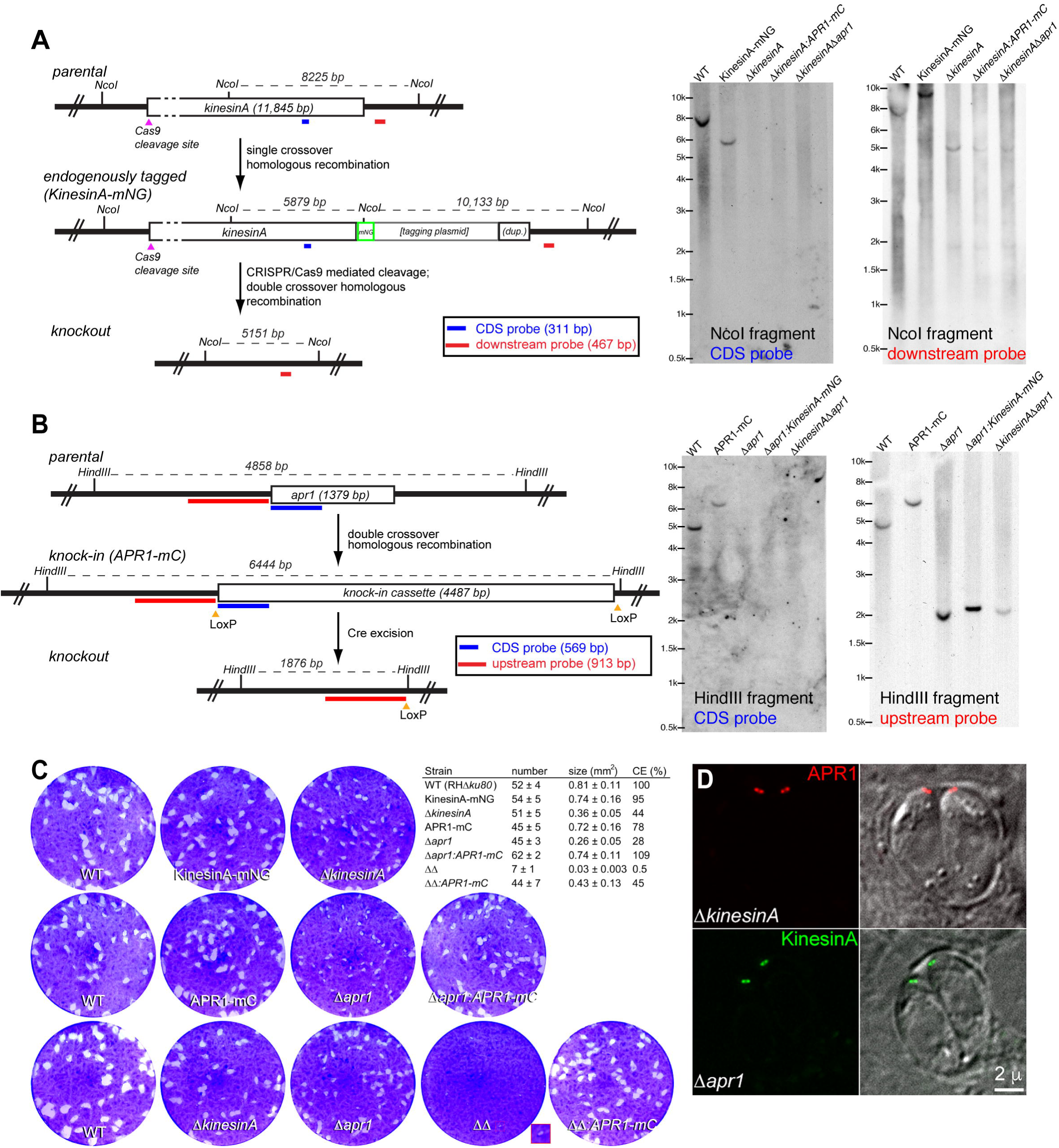
Figure 5. Generation of endogenously tagged *KinesinA-mNeonGreenFP, APR1-mCherryFP* knock-in, knockout and complemented parasites, and assessment of their plaquing efficiency. A. Left, schematic for generating endogenously tagged KinesinA and *ΔkinesinA* parasites and Southern blotting strategy. *RHΔku80* parasites (parental) were used to generate 3’ endogenously tagged *KinesinA-mNeonGreenFP* parasites [“endogenously tagged (KinesinA-mNG)”] via single crossover homologous recombination. mNG, *mNeonGreenFP* coding sequence; (dup.), *kinesinA* sequence partially duplicated as a result of the single crossover homologous recombination event. The endogenously tagged parasites were then transiently co-transfected with a plasmid expressing both Cas9 nuclease and a guide RNA specific for the 5’ end of the *kinesinA* genomic locus, and a repair plasmid containing the 5’ and 3’ flanking regions of *kinesinA* but lacking its coding sequence in between, to generate a knockout via double crossover homologous recombination. The mNeonGreenFP(-) parasite population was enriched using FACS prior to subcloning the knockout parasites. The positions of the Cas9 cleavage site, restriction sites, CDS probe (blue) and probe annealing downstream of the *kinesinA* genomic locus in the 3’ flanking region (red) used in Southern blotting analysis (right) and the corresponding DNA fragment sizes expected are shown. Right, Southern blotting analysis of the *kinesinA* locus in *RHAku80* (WT), *KinesinA-mNeonGreenFP* endogenously tagged *(KinesinA-mNG), ΔkinesinA, ΔkinesinA:APR1-mCherryFP (ΔkinesinA:APR1-mC)* and *ΔkinesinAΔapr1* parasites generated as described above. The expected parasite genomic DNA fragment sizes after *Nco*I digestion are, for the CDS probe, 8225 bp for the parental *(i.e.,* wild-type *kinesinA* locus), and 5879 bp for the endogenously tagged line. The expected DNA fragment sizes for the downstream probe are 8225 bp for the parental, 10,133 bp for the endogenously tagged, and 5151 bp for the knockout. B. Left, schematic for generating *APR1-mCherryFP* knock-in and *Δapr1 parasites* and Southern blotting strategy. *RHΔku80* parasites (parental) were used to generate knock-in *APR1-mCherryFP* knock-in parasites [“knock-in (APR1-mC)”] via double crossover homologous recombination. The knock-in parasites were then transiently transfected with a plasmid expressing Cre recombinase to excise the knock-in expression cassette between the two LoxP sites, and mCherryFP(-) parasites were enriched using FACS prior to subcloning the knockout parasites. The positions of the restriction sites, CDS probe (blue) and the probe annealing upstream of the *apr1* coding sequence (red) used in Southern blotting analysis and the corresponding DNA fragment sizes expected are shown. Right, Southern blotting analysis of the *apr1* locus in *RHΔku80* (WT), *APR1-mCherryFP* knock-in, *Δapr1, Δapr1:KinesinA-mNeonGreenFP (Δapr1:KinesinA-mNG)* and *ΔkinesinAΔapr1* parasites generated as described above. The expected parasite genomic DNA fragment sizes after *Hin*dIII digestion are, for the CDS probe, 4858 bp for the parental *(i.e.,* wild-type *apr1* locus), and 6444 bp for the knock-in. The expected DNA fragment sizes for the upstream probe are the same as for the CDS probe for the parental and knock-in, and 1876 bp for the knockout. C. Plaques formed by RH*Δku80* (WT), *KinesinA-mNeonGreenFP* endogenously tagged *(KinesinA-mNG)* and APR1-mCherry knock-in *(APR1-mC)* parasites, knockout *(ΔkinesinA, Δapr1, ΔkinesinAΔapr1*(*ΔΔ*)), and *Δapr1:.APR1-mCherryFP* (Δ*apr1:APR1-mC*) and *ΔkinesinAΔapr1:APR1-mCherryFP* (*ΔΔ*:*APR1*-mC) complemented parasite lines. HFF monolayers were infected with an equal number of each line of parasites, grown for 7 days at 37°C, and then fixed and stained with crystal violet. Host cells that remained intact (viable) absorbed the crystal violet staining, whereas regions of host cells lysed by the parasites (“plaques”) are clear. The box outlined in red is enlarged 2X in the inset to visualize the very small plaques formed by the *ΔkinesinAΔapr1* parasites. Inset table: quantification of the number and size of plaques (mean ± standard error) produced by the *T. gondii* lines, as measured in three independent biological replicates. The cytolytic efficiency (CE) is defined as total area of the host cell monolayer that has been lysed by a *T. gondii* line divided by the total area lysed by WT and expressed as a percentage. D. Localization of KinesinA-mNeonGreenFP (endogenous tagging) in live *Δapr1* parasites, and APR1-mCherryFP (knock-in) in live *ΔkinesinA* parasites.

The *apr1* knockout (*Δapr1*) parasites were generated by transient expression of Cre recombinase to excise the LoxP-flanked APR1-mCherryFP expression cassette as previously described (Heaslip *et al.,* 2010; Liu *et al.,* 2013; Liu *et al.,* 2016), and knock-in and knockout clones were confirmed by Southern blotting analysis (Figure 5B). The *Δapr1* parasites formed noticeably smaller plaques compared to either the wild-type or to the parental *APR1-mCherryFP* knock-in parasites (Figure 5C). Complementation with an ectopic copy of *APR1-mCherryFP* driven by the ~0.9 kb region immediately upstream of the *apr1* locus restored growth to wild-type levels.

In contrast to the moderate growth phenotype of the single knockout mutants, deletion of both *kinesinA* and *apr1* resulted in a severe defect in plaquing efficiency (Figure 5C). The double knockout line *(ΔkinesinAΔapr1* parasites) was generated using the same knock-in/knockout strategy as described above for the *Δapr1* parasites but with *ΔkinesinA* parasites as the parental line. The *ΔkinesinAΔapr1* parasites formed significantly fewer visible plaques seven days post-infection, and the plaques that could be observed were significantly smaller in diameter (Figure 5C). This growth defect could be partially compensated by re-introducing an ectopic copy of *APR1-mCherryFP* as described above. Comparison of the cytolytic efficiencies of the single and double knockout mutants relative to the wild type suggests that the combined loss of KinesinA and APR1 has a synergistic impact on the parasite lytic cycle. The cytolytic efficiency of the *ΔkinesinA* and *Δapr1* parasites is 44% and 28%, respectively, relative to the wild type. An additive effect of KinesinA and APR1 would predict the *ΔkinesinAΔapr1* parasites to have a relative cytolytic efficiency of ~12% (0.44*0.28), much higher than the observed 0.5%.

To determine whether KinesinA and APR1 are dependent on each other for localization to the apical polar ring, we examined the localization of a knock-in copy of *APR1-mCherryFP* in *ΔkinesinA* parasites and endogenously tagged KinesinA in *Δapr1* parasites. APR1-mCherryFP was localized to an apical ring in the *ΔkinesinA* parasite; similarly, endogenously tagged KinesinA still localized to an apical ring in the *Δapr1* parasites (Figure 5D). Their independent association with the apical polar ring and apparently synergistic mode of actions suggest that these two proteins might be involved in related, but differentially regulated functions.

### The loss of KinesinA and APR1 significantly reduces the efficiency of invasion and motility

Since the *ΔkinesinAΔapr1* parasites were defective in plaquing efficiency, we next characterized steps of the lytic cycle that could be influenced by the loss of *kinesinA* and *apr1.* As an obligate intracellular parasite, *T. gondii* needs to invade into host cells in order to proliferate. To examine how the loss of KinesinA and/or APR1 might affect the ability of parasites to invade, we performed a two-colour invasion assay that distinguishes invaded (intracellular) parasites from those that had not invaded (extracellular) by differential labelling of an antibody that recognizes an epitope of *Toxoplasma* surface antigen 1 (TgSAG1) (Carey *et al.,* 2004; Mital *et al.,* 2005). The loss of either *kinesinA* or *apr1* alone decreased invasion levels to roughly half that of wild-type parasites, and the loss of both resulted in an additive defect, reducing invasion levels to ~30% that of the wild type (Figure 6A and Table 1). Upon complementation of the double knockout mutant with a plasmid coding for *APR1-mCherryFP,* this invasion phenotype was restored back to a level comparable to the single *kinesinA* knockout.

**Figure 6.**
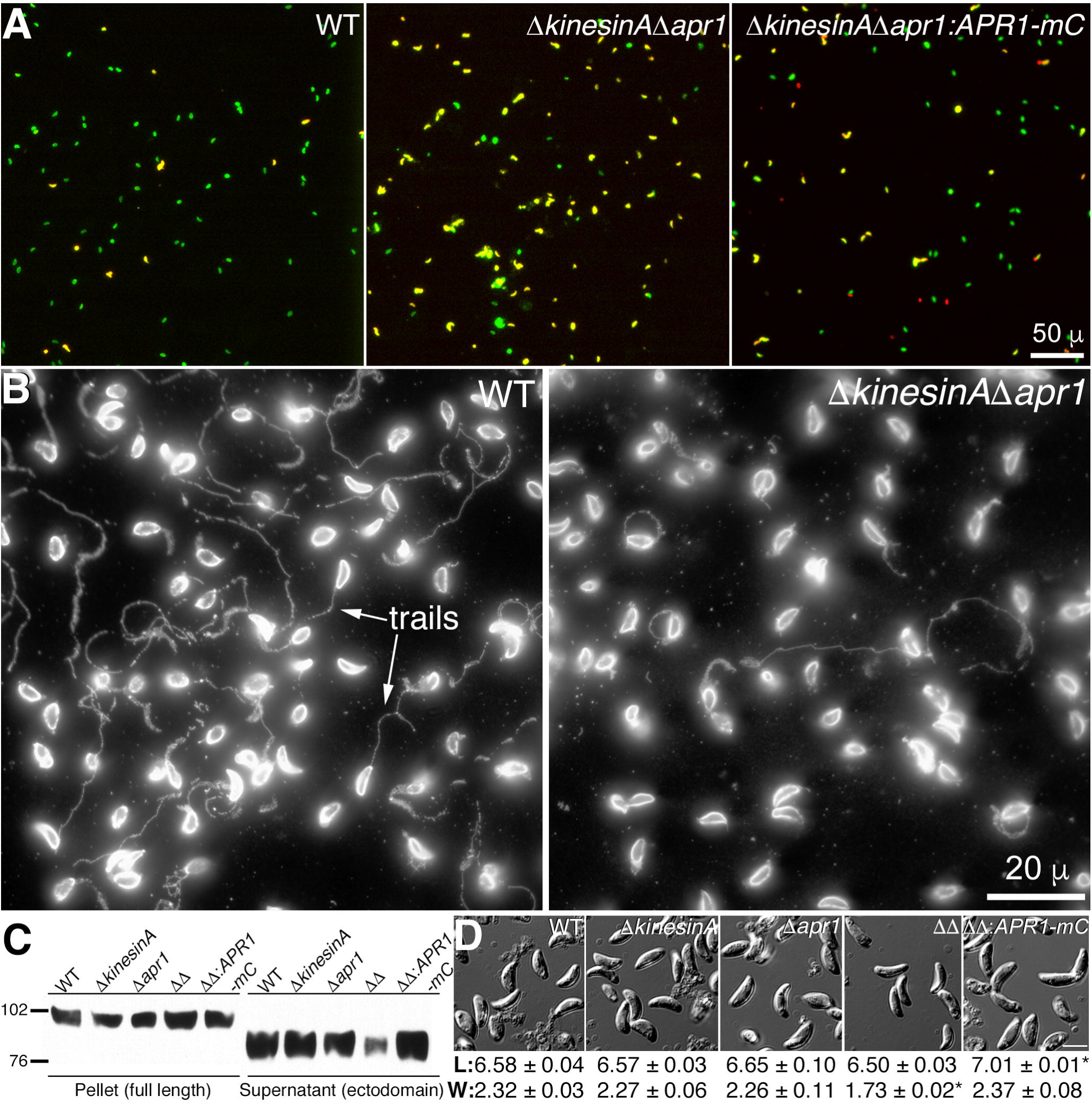
Parasite invasion, motility, and microneme secretion are impaired, and the parasite shape is altered when KinesinA and APR1 are absent. A. Representative wide-field epifluorescence images of invasion by the *T. gondii* lines indicated. Parasites that are intracellular *(i.e.,* have invaded) are labelled green, and parasites that did not invade are labelled both green and red *(i.e.* yellow). See also Table 1 for quantification of invasion assays. B. Wide-field epifluorescence images showing trail deposition of the RH*Δku80* (WT) or *ΔkinesinAΔaprl* parasites on coated dishes, as labelled by an antibody that recognizes *T. gondii* surface antigen 1 (TgSAGI). Images are representative of results from three independent biological replicates. C. The pellet and secreted fractions (supernatant) of parental, knockout and complemented parasite lines upon ethanol stimulation, as probed by antibodies against MIC2. Western blots shown are representative of three independent biological replicates. The pellet fraction contains the full-length and the supernatant contains the shed ectodomain of MIC2. The numbers on the left indicate molecular masses in kDa. The pellet and secreted fractions were cropped from different regions of the same western blot. D. The maximum length and width for live, extracellular RH*Δku8*0 (WT), *ΔkinesinA, Δapr1, ΔkinesinAΔapr1* (*ΔΔ*) and APR1-mCherryFP complemented *ΔkinesinAΔaprl (ΔΔ:APR1-mC)* parasites were measured using DIC imaging (representative set of images shown). The experiment was performed in triplicate; values are the mean ± standard error μm) from 100 parasites per parasite line in each experiment. (unpaired Student’s t-test, * p < 0.0001 compared to the wild type). Length (L) and width (W) are in measured in μm. Scale bar, 5 μm.

**Table 1.** Quantification of invasion (mean number of intracellular parasites ± standard error) by the *T. gondii* lines indicated. The number of intracellular parasites per field was counted in ten fields per strain, in each of three independent biological replicates. P-values from unpaired, two-tailed Student’s t-tests are indicated on the right.

| Strain | Invasion Assay |  | P-value |  |  |  |  |
| --- | --- | --- | --- | --- | --- | --- | --- |
| | # invaded | % WT | RHΔku80 | ΔkinesinA | Δapr1 | ΔkinesinAΔapr1 | ΔkinesinAΔapr1:<br>APR1-mC |
| RHΔku80 (WT) | 256 ± 29 | 100 | | 0.028 | 0.022 | 0.0038 | 0.0093 |
| ΔkinesinA | 138 ± 19 | 54 | | | 0.99 | 0.035 | 0.34 |
| Δapr1 | 139 ± 14 | 54 | | | | 0.014 | 0.21 |
| ΔkinesinAΔapr1 | 75 ± 6 | 29 | | | | | 0.0037 |
| ΔkinesinAΔapr1:<br>APR1-mC | 118 ± 3 | 46 | | | | | |

Invasion and parasite motility are tightly linked, and we therefore next examined the *ΔkinesinAΔapr1* parasites for motility defects in a trail deposition assay. Parasites form “trails” rich in lipids and proteins such as TgSAG1 as they glide upon a substrate-coated surface (Hakansson *et al.,* 1999; Carey *et al.,* 2004). The deposited trails can then be visualized by indirect immunofluorescence, and evaluated in terms of the number, length and type to correlate to parasite motility levels. Parasites undergoing circular gliding deposit “rings” of SAG1-positive trails, whereas parasites undergoing helical gliding leave behind “scalloped” trails thought to be indicative of productive, directional movement. There was a qualitative difference in the number of SAG1-positive trails that the *ΔkinesinAΔapr1* parasites deposited compared to the parental parasites, and within this motile fraction of the population, there appeared to be a higher proportion of circular trails and short trails *(i.e.,* less than one parasite length), suggesting the *ΔkinesinAΔapr1* parasites have a motility defect (Figure 6B). Interestingly, some *ΔkinesinAΔapr1* parasites were able to deposit trails during helical gliding that resembled those formed by wild-type parasites in length and shape. One possible explanation for this phenotype is that the efficiency of motility initiation in the *AkinesinAAaprl* parasites is poor compared to the wild type, but once initiation has occurred, then the parasite can accomplish directional movement.

### The loss of KinesinA and APR1 significantly reduces the induced secretion of adhesins

Secretion of proteins from the micronemes is an important step for motility initiation as well as for invasion into host cells. For instance, transmembrane proteins including micronemal protein 2 (MIC2) and MIC2-associated protein (M2AP) are secreted onto the plasma membrane of the parasite (Carruthers and Sibley, 1997; Carruthers *et al.,* 1999a; Huynh *et al.,* 2003; Huynh and Carruthers, 2006), where they are proposed to link the actomyosin machinery to the host cell surface, such that activity of the motor complex generates productive parasite movement (Opitz and Soldati, 2002; Sibley, 2010). Consistent with this view, the knockdown of *mic2* expression levels results in a greater proportion of non-motile parasites and parasites unable to sustain productive motility; these parasites also have a significant defect in host-cell invasion (Huynh and Carruthers, 2006). Secretion of micronemal proteins can be triggered upon treatment of parasites with ethanol (Carruthers *et al.,* 1999 b; Lovett *et al.,* 2002; Lourido and Moreno, 2015). These proteins are cleaved by transmembrane proteases after secretion from the apical end of the parasite, and the amount of protein released in the soluble fraction is used as an indicator of secretory activity from the micronemes. Using this assay, we observed a pronounced decrease in ethanol-induced secretion of MIC2 from the *ΔkinesinAΔapr1* parasites compared to the wild-type and single knockout lines, and secretion was restored with ectopic expression of *APR1-mCherryFP* (Figure 6C). The impaired secretion of this adhesin likely contributes to the decrease in invasion and motility of parasites lacking KinesinA and APR1.

### The loss of KinesinA and APR1 alters parasite shape and significantly reduces the mechanical stability of the apical polar ring

It was shown previously that parasites lacking a cytoskeleton-associated component can have an altered shape (Egarter *et al.,* 2014; Leung *et al.,* 2014) as well as diminished motility (Leung *et al.,* 2014). Indeed, the extracellular *ΔkinesinAΔapr1* parasites had a different morphology from that of the wild-type and the single knockout parasites. While there was no difference in the mean length, the double knockout parasites were significantly thinner than the wild-type or single knockout parasites. Stable expression of *APR1-mCherryFp* in the double knockout parasites was sufficient to restore the width to wild-type levels (Figure 6D). There was also a modest but statistically significant increase in mean parasite body length, although the length:width ratio was not significantly different from the wild-type parasite (unpaired Student’s t-test, p-value of 0.29).

The difference in parasite shape suggested there were changes in the underlying cytoskeletal architecture of the *ΔkinesinAΔapr1* parasites. We therefore investigated how the loss of KinesinA and APR1 at the apical polar ring might affect the organization and stability of the cortical microtubule array. Specifically, we addressed the following questions: 1) Can parasites lacking KinesinA, APR1 or both, generate an array of 22 cortical microtubules? 2) If they can, does the organization of the cortical microtubules differ from that of the wild type? 3) Are KinesinA and APR1 involved in maintaining the mechanical stability of the apical polar ring and/or the cortical microtubules?

To address these questions, we first examined sonicated extracellular *ΔkinesinAΔapr1* parasites in which the cortical microtubules were spread out radially on the coverslip, a convenient arrangement that allowed us to count the number of cortical microtubules by immunofluorescence and anti-tubulin antibodies. We found that *ΔkinesinAΔapr1* parasites were capable of making 22 cortical microtubules (Figure 7A), which indicates that neither KinesinA nor APR1 is essential in templating the 22 cortical microtubule array. However, examination of the cortical microtubules in live parasites expressing mEmeraldFP-Tg-α1- tubulin showed that even though the arrangement and spacing of the cortical microtubules in some *ΔkinesinAΔapr1* parasites appear to resemble that of the wild type, the microtubule array appears abnormal in many, with groups of two or three cortical microtubules detached from the apical end, leaving gaps in the array (Figure 7B). Immunofluorescence labeling by tubulin antibody shows similar results *(c.f.* Figure 9). Furthermore, RNG1 – a marker for the apical polar ring - endogenously tagged with mCherryFP in the *ΔkinesinAΔapr1* parasites did not form a complete ring as was seen in 100% of the wild-type parasites (Tran *et al.,* 2010) (Figure 7B and Figure S2). Instead, in 100% of the *ΔkinesinAΔapr1* parasites, the RNG1-mCherryFP signal was localized to multiple discrete dots, regardless of whether or not the disorganization of the microtubule array was apparent (Figure 7B, *ΔkinesinAΔapr1).* The RNG1-mCherryFP spots were often found at the roots of groups of microtubules that had detached from the apical end. This suggests that the structure of the apical polar ring is compromised in live *ΔkinesinAΔapr1* parasites, and detachment of the cortical microtubules from the apex is likely a result of this structural defect. The extent of detachment *(i.e.* whether or not an obvious gap in the array can be observed) appears to be stochastic, since it differs between parasites in the same vacuole *(e.g.* Figure 7B, bottom row).

**Figure 7.**
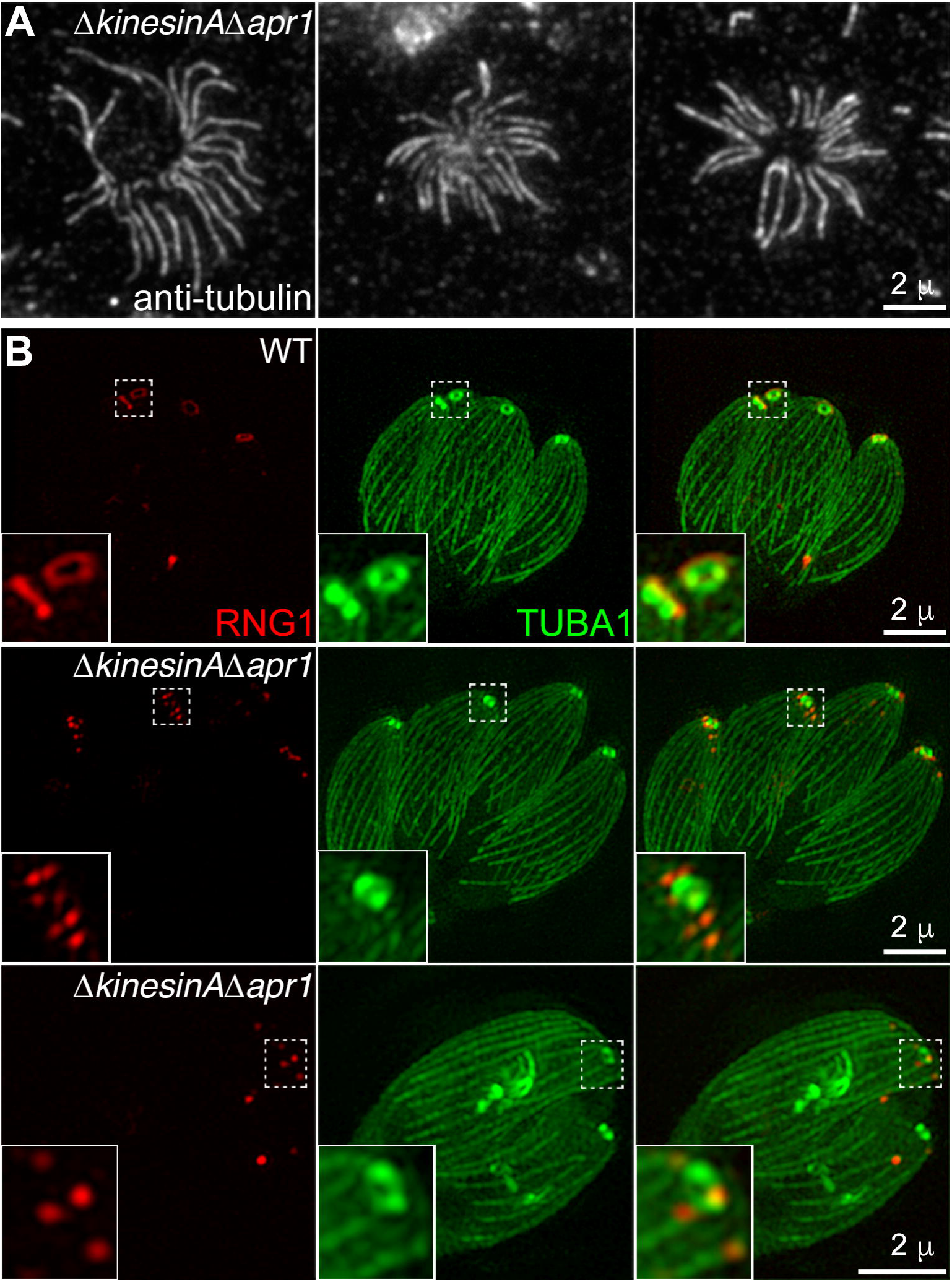
*ΔkinesinAΔaprl* parasites are still capable of making 22 cortical microtubules, but the loss of KinesinA and ApR1 impairs the stability of the apical polar ring. A. Wide-field epifluorescence images of extracellular *ΔkinesinAΔaprl* parasites that were briefly sonicated, and labelled with anti-tubulin antibodies to count the number of splayed cortical microtubules. Shown are three examples, each with 22 cortical microtubules. B. 3D-SIM projections of live, intracellular *RHΔku80* (WT) and *ΔkinesinAΔapr1* parasites stably expressing RNG1 endogenously tagged with mCherryFP (RNG1, red), and transiently expressing mEmeraldFP-α1-tubulin (TUBA1, green) from a *T. gondii* tubulin promoter. Insets are shown at 3X.

To further understand the structural role(s) of KinesinA and APR1, we tested the stability of the apical polar ring in the single and double knockout mutants when extracted by detergent. In wild-type parasites, the apical polar ring is highly resistant to detergent extraction ((Russell and Burns, 1984; Nichols and Chiappino, 1987; Hu *et al.,* 2002; Tran *et al.,* 2010), Figure 8A). The structure of the apical polar ring in the parental line stably expressing KinesinA-mNeonGreenFP was indistinguishable from wild-type parasites (data not shown), indicating that the initial endogenous tagging of KinesinA did not significantly alter the stability of the apical polar ring, as judged by this assay. When *kinesinA* was deleted from the genome, the apical polar ring was no longer detectable by transmission electron microscopy and negative staining with phosphotungstic acid, upon brief extraction (3 min.) with 0.5% (v/v) Triton X-100 (TX-100), a non-ionic detergent (Figure 8A and Figure S3). This was observed in any parasite line that lacked *kinesinA (i.e. ΔkinesinA, ΔkinesinAΔapr1* and *ΔkinesinAΔapr1* parasites complemented with APR1-mCherryFP). In some instances, the roots of the cortical microtubules abut the base of an electron-lucent region (Figure 8A, extended *ΔkinesinA* and extended *ΔkinesinAΔapr1:APR1-mC)* where the apical polar ring would have been visible in wild-type parasites, while in others a thin line of electron-dense material is present between the roots (Figure 8A, extended *ΔkinesinAΔapr1,* black arrowheads). In contrast, deletion of *apr1* did not affect the resistance of the apical polar ring to TX-100 extraction, but resulted in a higher incidence of partial or complete detachment of the conoid from the parasite compared to the wild-type *(RHΔku80), APR1-mCherryFP* knock-in or complemented *Δapr1* parasites [RH*Δ*ku80, 6% conoids partially or fully detached (n=35); *APR1-mCherryFP* knock-in, 0% (n=20); *Δapr1,* 64% (n=25); complemented *Δapr1,* 12% (n=25); and *ΔkinesinAΔapr1,* 57% (n=28)] (Figure 8A and Figures S3-S7). The defects in the *kinesinA* and *apr1* mutants are remarkably specific, as the other structures in the cytoskeletal apical complex, including the conoid, preconoidal rings, and intra-conoid microtubules appear to be unperturbed. The cortical microtubules, even though apparently anchorless in lines lacking *kinesinA,* also remain stable. Conversely, those other cytoskeletal structures can be dismantled specifically without affecting the apical polar ring. The ring remains intact when the cortical and intra-conoid microtubules are destabilized due to the loss of three associated proteins *(Δtlap2Δspm1Δtlap3,* Figure 8A, (Liu *et al.,* 2016; Nagayasu *et al.,* 2016)). It is also unaffected by the disruption of the conoid structure when a component of the conoid fibers is removed (Nagayasu *et al.,* 2016).

**Figure 8.**
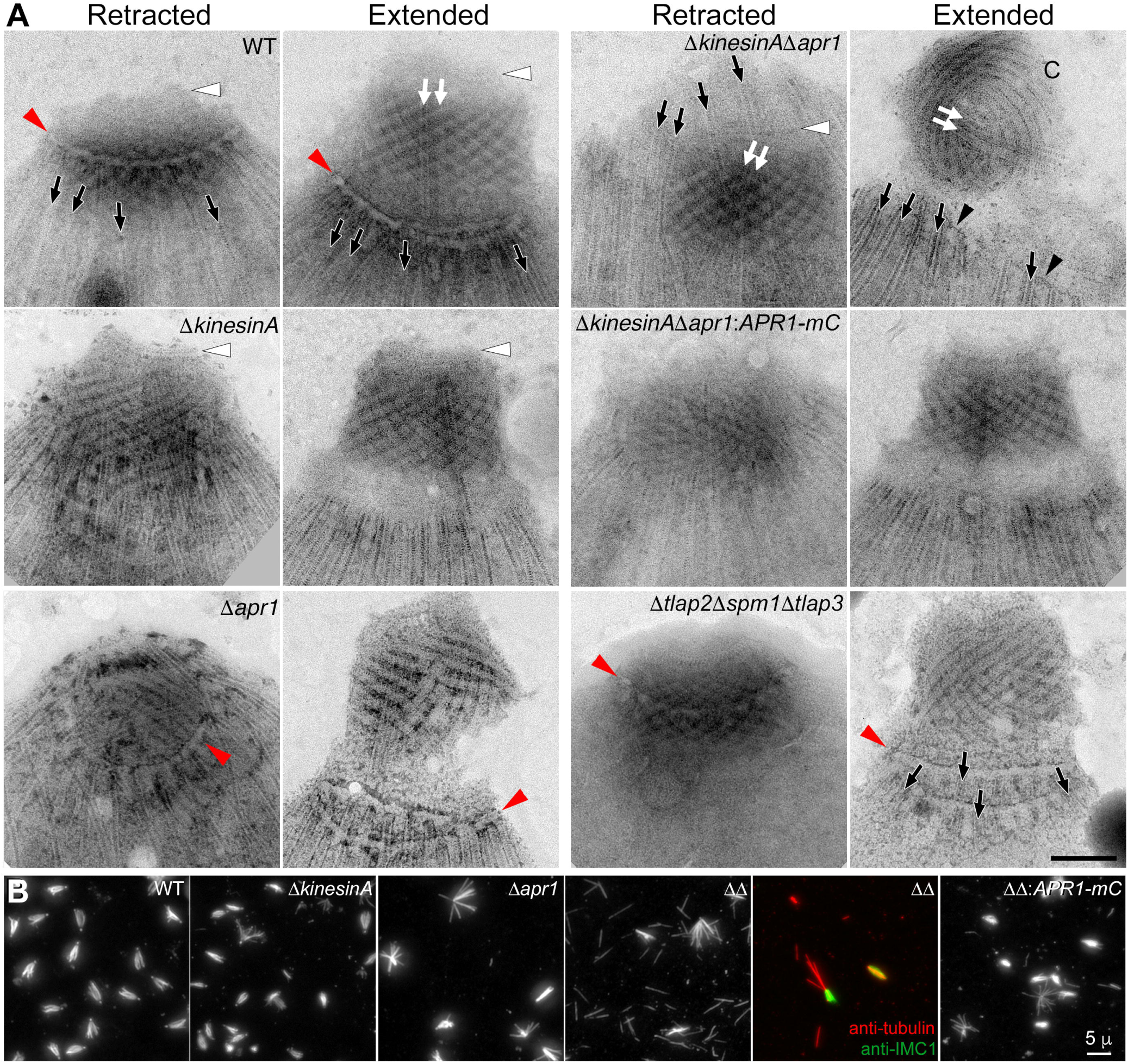
The apical polar ring is unstable in parasites lacking KinesinA when extracted by detergent. A. Transmission electron micrographs of negatively stained, whole mount Triton X-100- extracted *RHΔku80* (WT), Δ*kinesinA,* Δ*apr1,* Δ*kinesinA*Δ*apr1* and Δ*kinesinA*Δ*apr1* complemented with APR1-mCherryFP (Δ*kinesinA*Δ*apr1:APR1-mC),* and Δ*tlap2*Δ*spm1*Δ*tlap3* parasites. See also Figures S3-S7 for additional EM images. Images shown are representative of at least two independent biological replicates. Retracted, parasites with a retracted conoid; extended, parasites with an extended conoid. Red arrowheads: the apical polar ring; white arrowheads, preconoidal rings; black arrows, cortical microtubules; white arrows, intra-conoid microtubules; black arrowheads, a thin line of electron-dense material between the roots of the cortical microtubules; C, conoid. Scale bar, 200 nm. B. Wide-field epifluorescence images of extracellular RH*Δku80* (WT), Δ*kinesinA,* Δ*apr1,* Δ*kinesinA*Δ*apr1* (ΔΔ) and Δ*kinesinA*Δ*apr1* complemented with APR1-mCherryFP (ΔΔ*-.APRI-mC)* parasites, extracted with 10 mM sodium deoxycholate, and subsequently fixed with 3.7% (v/v) formaldehyde and labelled with a mixture of mouse anti-α- and β-tubulin antibodies. Extracted Δ*kinesinA*Δ*apr1* parasites were also labelled with rabbit anti-IMC1 antibody (green) to show that the dissociated cortical microtubules (red) appear to be free of parasite cortex. Images shown are representative of three independent biological replicates. Scale bar, 5 μm.

In some TX-100 extracted *ΔkinesinAΔapr1* parasites, the array of cortical microtubules is splayed and no longer converges at the parasite apex at the EM level (Figure 8A and Figure S3), suggesting an overall weakened association. Indeed, the cortical microtubules of *ΔkinesinAΔapr1* parasites can be completely dissociated from the parasite upon extraction with an anionic detergent, sodium deoxycholate, resulting in free microtubules scattered across the field (Figure 8B). The microtubules that do remain associated with the extracted parasite cortex (labelled by an anti-IMC1 antibody) sometimes collapse into a bundle within the parasite. In contrast, the cortical microtubules in most wild-type parasites, the single knockout mutants and the APR1-mCherryFP-complemented double knockout mutant remain clustered at the apical end of the parasites. This further confirms that KinesinA and APR1 play related but independent structural roles.

### The perturbation in the microtubule cytoskeleton correlates with a change in the distribution of micronemal vesicles

The perturbation in the organization of the cortical microtubules is linked with a change in the distribution of micronemal vesicles within the parasite. In intracellular wild-type parasites, these vesicles – labelled by anti-MIC2 antibodies – are predominantly located in the apical half of the parasite, and appear to be organized into tracks that co-align with the cortical microtubules labelled by anti-Tg-β2-tubulin antibodies (Figure 9A). Large gaps in the microtubule array in *ΔkinesinAΔapr1* parasites correlate with the absence of MIC2-containing vesicles (Figure 9B). Furthermore, when the cortical microtubules are partially polymerized in a parasite line *(Δtlap2Δspm1Δtlap3)* lacking three microtubule-associated proteins (Liu *et al.,* 2016), the micronemal vesicles are concentrated in the region where the remnant cortical microtubules are (Figure 9C, “37°C”). In *Δtlap2Δspm1Δtlap3* parasites where the cortical microtubules are mostly destabilized after incubation at low temperatures (Liu *et al.,* 2016), the micronemal vesicles are no longer concentrated in the apical portion of the parasite, even though some vesicles still appear to be aligned in tracks (Figure 9C, “cold-treated”, white arrows). Incubating wild-type parasites in the same low temperature conditions did not affect the distribution of MIC2-containing vesicles, indicating the cold treatment itself did not cause the altered vesicle distribution (data not shown).

**Figure 9.**
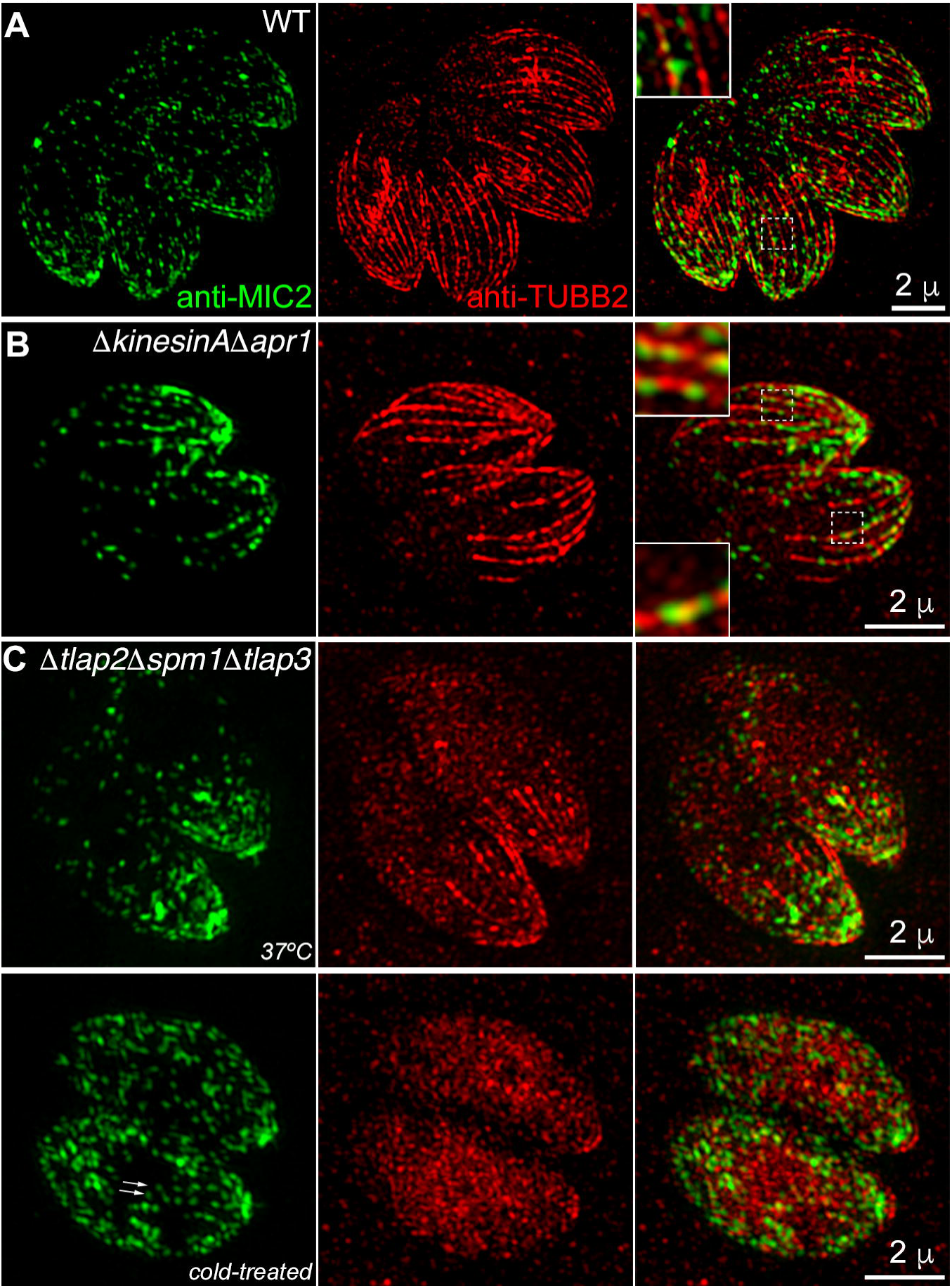
**Figure 9.** MIC2-containing microneme vesicles are aligned with cortical microtubules, and their distribution is altered in parasites whose cortical microtubule organization has been perturbed. 3D-SIM half-volume projections of fixed, intracellular (A) *RHΔku80* (WT), (B) Δ*kinesinA*Δ*apr1*, and (C) Δ*tlap2*Δ*spm1* Δ*tlap3* parasites that had been either incubated in low temperature conditions (“cold-treated”) or not (“37°C”) prior to fixation and labelling with a rabbit anti-Tg-β2- tubulin (TUBB2) antibody (Morrissette and Sibley, 2002b) (red) and a mouse anti-MIC2 antibody (green) (Carruthers *et al.,* 2000). Arrows indicate MIC2 signal aligned in tracks in cold-treated Δ*tlap2*Δ*spm1*Δ*tlap3* parasites. Note that the anti-Tg-β2-tubulin antibody does not label the conoid under these conditions, presumably due to antigen accessibility issues (see also (Liu *et al.,* 2016)). Half-volume rather than full-volume projections are presented for optimal visualization of the localization of MIC2-containing vesicles, relative to the cortical microtubules. Insets are shown at 3X. Scale bar, 2 μm.

## DISCUSSION

In this work, we have identified and characterized two new components - KinesinA and APR1 - of the apical polar ring, a putative organizing center for the 22 cortical microtubules in the human parasite, *Toxoplasma gondii.* The well-defined structure of the apical polar ring and its connection with the microtubules is a useful model in general for understanding how microtubules can be stably anchored. We found that KinesinA and APR1 are important for anchoring the cortical microtubules by maintaining the stability of the apical polar ring and for efficient secretion of adhesins from the micronemes. The combined loss of these two proteins results in a dramatic retardation in the lytic cycle of the parasite. Importantly, the stability of the apical polar ring can be separated from that of associated cytoskeletal structures, *i.e.* the conoid, preconoidal rings, intra-conoid microtubules and cortical microtubules. This highlights the modularity of the cytoskeletal framework and demonstrates that it is possible to explore the functions of these structures both individually and as an ensemble.

### The construction and stability of the apical polar ring

KinesinA and APR1 are recruited to the apical polar ring during the early and late stages of daughter construction, respectively. Although KinesinA is predicted to contain a kinesin-like motor domain, the signal of fluorescently-tagged KinesinA is always concentrated in a disc-like structure at the apical end of the parasite, and is not observed along the cortical microtubules at any stage during the cell cycle. This suggests that the recruitment of KinesinA to the daughter apical polar ring is direct rather than a result of convergence of the KinesinA molecules to the apical end via transport along the cortical microtubules, and that KinesinA probably does not act like a conventional motor, such as Pkl1 in *Schizosaccharomyces pombe* that transports proteins to the spindle pole body and anchors the microtubules (Pidoux *et al.,* 1996; Yukawa *et al.,* 2015). However, it remains possible that KinesinA might function as a motor during a very short timeframe that so far has escaped our attention. The ring-like KinesinA signal in nascent daughters suggests that the overall architecture and initial level of stability of the apical polar ring is established early on, but the addition of components such as APR1 is needed to reinforce the stability of the apical polar ring itself and/or its association with other structures such as the conoid. This multi-level fortification is reflected in the different structural defects observed in *kinesinA* and *apr1* knockout parasites upon detergent extraction. In the future, it will be of interest to know at what stage of daughter replication the fragmentation of the apical polar ring initially occurs in the *ΔkinesinAΔapr1* parasite. It would also be informative to investigate how other components of the apical polar ring, such as RNG1 (recruited at a late stage of daughter formation to the apical polar ring) and RNG2 (recruited during early stages) (Tran *et al.,* 2010; Katris *et al.,* 2014), might contribute to the assembly and/or biogenesis of this structure, and how their functions could be affected in the absence of KinesinA, or APR1, or both.

### Stability of the apical polar ring is not tightly coupled with its potential templating function

The regularly spaced 22 cortical microtubules are anchored to the apical polar ring ((Russell and Burns, 1984; Nichols and Chiappino, 1987), Figure 1). This observation has long supported the hypothesis that the apical polar ring is an organizing center and provides a template for the biogenesis of the cortical microtubules. Although the apical polar ring is destabilized, *ΔkinesinAΔapr1* parasites are capable of generating 22 cortical microtubules *(c.f.* Figure 7A). This indicates that the stability of the apical polar ring can be separated from its potential function in the nucleation and templating of cortical microtubules. Gamma-tubulin, the nucleator in canonical microtubule organizing centers, is detected by immunofluorescence in the centrosome in *T. gondii,* but not the apical polar ring (Suvorova *et al.,* 2015), suggesting that an unusual mechanism is responsible for nucleating the cortical microtubules. Besides the nucleating factors, we also do not know what the “rulers” in the apical polar ring are that establish the 22-fold rotational symmetry, or how the nucleation and spatial arrangement of the cortical microtubules are coordinated. To this end, identification of additional components of the apical polar ring is needed, for which the two components that had been characterized previously (RNG1 and RNG2, (Tran *et al.,* 2010; Katris *et al.,* 2014)) as well as the two newly identified here (KinesinA and APR1) will be useful probes.

### The role of the apical polar ring and microtubule cytoskeleton in the parasite lytic cycle

The structural defects in the apical polar ring are linked to impaired parasite motility and invasion, and a severe overall retardation in the parasite lytic cycle. This can be partially explained by a reduction in secretion of the adhesin MIC2 accompanying the loss of KinesinA and APR1, as MIC2 is important for the adhesion of the parasite to its environment during motility and invasion. Although it has been shown that, as with regulated secretory processes in other systems, an increase in free cytoplasmic [Ca^2+^] in the parasite enhances secretion (Carruthers and Sibley, 1999; Lovett *et al.,* 2002), the structural framework that supports the polarized transport of micronemal vesicles to and out the parasite apex remains unknown. There are several possible explanations for the defect in microneme secretion observed in our mutants. Structural perturbation of the apical polar ring itself could disrupt exit of the vesicles through the apical complex, or the loss of KinesinA and APR1 could interfere with functions of other apical polar ring components, such as RNG2, which affects microneme secretion through the cGMP-Ca^2^+ signalling pathway (Katris *et al.,* 2014). Alternatively, the secretion defect might be coupled with the change in the organization of the cortical microtubules, which appear to be well-aligned with tracks of MIC2-containing vesicles (Figure 9). For instance, if the cortical microtubules are used to transport the micronemal vesicles, then the disorganization of these polymers could affect transport efficiency. On the other hand, the cortical microtubules might be used instead as temporary storage sites for the vesicles, since the heavy coating of cortical microtubules by stably associated proteins (Tran *et al.,* 2012; Liu *et al.,* 2013; Liu *et al.,* 2016) might impede efficient movement of motor proteins along these polymers. Future studies that explore secretion in live parasites are needed to test some of these possibilities, for which the set of mutants with different defects in the organization and stability of cortical microtubules (this study, and (Liu *et al.,* 2016)) will be useful.

The specific structural defect in the apical polar ring impacts multiple aspects of the physiology of the parasite. In addition to the reduction in MIC2 secretion, other phenotypes, such as the change in parasite shape, might also contribute to the significant defect in motility and invasion in the parasites lacking KinesinA and APR1. Furthermore, weakening of the apical polar ring and its association with the conoid might directly affect the ability of the parasite to overcome mechanical barriers as it penetrates the host cell during invasion.

In summary, this work has revealed certain structural requirements for the stability of the apical polar ring as well as a link between the microtubule cytoskeleton and parasite secretion. In the future, the components of the apical polar ring identified in this and previous studies will serve as valuable probes to determine the composition of the apical polar ring comprehensively, for addressing if and how this structure dictates the number and spacing of the cortical microtubules. Finally, structural perturbations of the apical polar ring make it possible to isolate the cortical microtubules from the membrane pellicle, which paves the way for examining their biophysical properties *in vitro,* including capping status, polymerization kinetics, and modulation of motor activity.

## MATERIALS AND METHODS

### *T. gondii,* host cell cultures, and parasite transfection

Tachyzoite *T. gondii* parasites were maintained by serial passage in confluent human foreskin fibroblast (HFFs, ATCC# SCRC-1041, Rockville, MD) monolayers in Dulbecco’s Modified Eagle’s Medium (DMEM, Life Technologies-Gibco, Cat# 10569-010), supplemented with 1% (v/v) heat-inactivated cosmic calf serum (Hyclone, Cat# SH30087.3) as previously described (Roos *et al.,* 1994; Leung *et al.,* 2014; Liu *et al.,* 2016). African Green Monkey renal epithelial cells (BS-C-1, ATCC# CCL-26) and rat aortic smooth muscle cells (A7r5, ATCC# CRL-1444) were cultured in the same manner as HFFs. *T. gondii* transfections were carried out as previously described (Liu *et al.,* 2013).

### Cloning of plasmids

Primers used in cloning DNA fragments and for sequencing are listed in Table S1. Genomic DNA (gDNA) fragments were amplified using gDNA template prepared from RHΔ*hx* or RHΔ*ku80*Δ*hx* (“RΗΔ*Ku80*”) parasites ((Fox *et al.,* 2009; Huynh and Carruthers, 2009); a kind gift from Dr. Vern Carruthers, University of Michigan, Ann Arbor, MI) using the Wizard Genomic DNA Purification Kit (Cat# A1120, Promega, Madison, WI) according to the manufacturer’s instructions. Similarly, coding sequences (CDS) were amplified using *T. gondii* complementary DNA (cDNA). All DNA fragments generated by PCR were confirmed by sequencing.

pTKO2_V-TgAPR1-mCherryFP, for generation of *APR1-mCherryFP* knock-in parasites: This plasmid was constructed in the pTKO2_V plasmid backbone designed for replacement of genes in *T. gondii* by homologous recombination. pTKO2_V was derived from pTKO2_II (Heaslip *et al.,* 2010) by replacing the coding sequence for *GFP* on the backbone with that for *EGFP* mutagenized to remove several restriction sites. Otherwise, pTKO2_V and pTKO2_II are identical. To generate pTKO2_V-TgAPR1-mCherryFP, three PCR fragments were generated and ligated with the pTKO2_V vector backbone using the corresponding sites. The 5’UTR of *apr1*(TgGT 1_315510) was amplified using primers S1 and AS1, and ligated via the *Not*I and EcoRI sites. The 3’UTR was amplified using primers S2 and AS2, and ligated via the HindIII and ApaI sites. The *APR1-mCherryFP* CDS was amplified using primers S3 and AS3 and ligated via the *Bgl*II and RsrII sites. The template for the *APR1-mCherryFP* CDS PCR was generated as follows: the coding sequence for *apr1* was first amplified using primers S4 and AS4, with cDNA template prepared from RHΔ*hx* parasites. The CDS was then ligated with the intermediate vector pdeg02 (see below) via FseI and *Sgr*DI to generate an *APR1-mCherryFP* fusion gene.

pdeg02 (the intermediate vector for generating pTKO2_V-TgAPR1-mCherryFP): The 7879 bp fragment liberated by *Xho*I from plasmid ptub_EGFP-mCherryFP (map and sequence in Figure S8) was religated with itself to give plasmid ptub_no-cat. The 6409 bp fragment released from ptub_no-cat by EcoRI and *Afl*II was ligated with a 407 bp *MfeI-Afl*III fragment derived from a synthesized 428 bp piece to give plasmid pdeg01. The synthesized piece consisted of (5’ to 3’) a multiple cloning site (MCS) with recognition sites for 15 restriction enzymes, beginning with *Mfe*I and ending with *Mre*I*-EcoR*I, followed by the 321 bp coding sequence for *FKBP(L106P)* (Banaszynski *et al.,* 2006), and ending with a stop codon plus an *Afl*II site. Plasmid pdeg02 was prepared from pdeg01 that was digested with *Mre*I and *Eco*RI, ligated with an *mCherryFP* coding sequence prepared by PCR from plasmid pRSET-B_mCherryFP (Shaner *et al.,* 2004) with primers S5 and AS5.

ptubA1-APR1-mCherryFP and ptubA1-APR1-EGFP: The coding sequences for *APR1-mCherryFP* or *APR1-EGFP* with a linker sequence coding for SSGLRS in between *apr1* and *mCherryFP* or *EGFP* were cloned into the *Nhe*I and *Afl*II sites in the same backbone as the plasmid ptub_EGFP-mCherryFP (map and sequence in Figure S8). The coding sequence of *apr1* contains the ATG start codon and is flanked by *NheI* on the 5’ end and *Bgl*II on the 3’ end. The coding sequences for *mCherryFP* and *EGFP* lack the ATG start codon and are flanked by *Bgl*II on the 5’ end and *Afl*II on the 3’ end.

ptubA1_mCherryFP-TgTUBA1: The coding sequences for *mCherryFP-TgTUBAI* with a linker sequence coding for SGLRS between *mCherryFP* and *α1-tubulin* were cloned into the *Nhe*I and *Afl*II sites in the same backbone as the plasmid ptub_EGFP-mCherryFP (map and sequence in Figure S8). The coding sequence of *mCherryFP* contains the ATG start codon and is flanked by *Nhe*I on the 5’ end and *Bgl*II on the 3’ end. The coding sequence for *α1- tubulin* lacks the ATG start codon and is flanked by *Bgl*II on the 5’ end and *Afl*II on the 3’ end.

ptubA1_mEmeraldFP-TgTUBA1: The *mEmeraldFP* coding sequence was amplified using primers S6 and AS6, using pUC57_mEmeraldFP-TgTrxL1 (Liu *et al.,* 2013) as the template, and ligated with ptubA1_mCherryFP-TgTUBA1 via *Nhe*I and *Bgl*II to generate the plasmid ptubA1_mEmeraldFP-TgTUBA1.

ptubA1g_mAppleFP-TgTUBB1: ptubA1g_mAppleFP-TgTUBB1 was generated with a three-component assembly using the NEBuilder HiFi DNA Assembly kit (NEB# E2621S) according to the manufacturer’s instructions. The vector backbone was prepared by removing the 12 bp stuffer sequence in ptubg (Nagayasu *et al.,* 2016) using *Nhe*I and *Afl*II. The *mAppleFP* coding sequence was amplified from plasmid pmAppleFP_C1 ((Kremers *et al.,* 2009), a kind gift from Richard Day, Indiana University School of Medicine, Indianapolis, IN) using primers S7 and AS7. The coding sequence for *β1-tubulin* was amplified from ptub-EGFP-TUBB1 (Nagayasu *et al.,* 2016) using primers S8 and AS8.

pTKO2_C-replace_TgKinesinA_mNeonGreenFP, for 3’ endogenous tagging of TgKinesinA with mNeonGreenFP: First, the gDNA region spanning the 3’ end of Intron 9 to Exon 10 of *kinesinA* (TgGT1_267370) was amplified using primers S9 and AS9, and ligated with pTKO2_II_3_MyoF_mEmeraldFP (Heaslip *et al.,* 2016) via EcoRI and *BglII* to generate the plasmid pTKO2_C-replace_TgKinesinA_Step1, which lacks a stop codon at the 3’ end of Exon 10. Next, the *mNeonGreenFP* coding sequence was amplified using primers S10 and AS10 with the plasmid pmNeonGreenFP-N1 as a template ((Shaner *et al.,* 2013), a kind gift from Richard Day, Indiana University School of Medicine, Indianapolis, IN), and ligated with pTKO2_C-replace_TgKinesinA_Step1 via *Bgl*II and *Afl*II. This yielded the plasmid pTKO2_C-replace_TgKinesinA_NeonGreen_Step1. The genomic fragment spanning the 3’ end of *kinesinA* Intron 8, Exon 9 and 5’ end of Intron 9 was amplified using primers S11 and AS11, and ligated with pTKO2_C-replace_TgKinesinA_NeonGreen_Step1 via *Avr*II and *Not*I to generate the 3’ endogenous tagging plasmid pTKO2_C-replace_TgKinesinA_mNeonGreenFP.

pTKO2_C-replace_TgKinesinB_mNeonGreenFP, for 3’ endogenous tagging of TgKinesinB with mNeonGreenFP: First, the gDNA region spanning the 3’ end of Intron 21 to the 3’ end of Exon 22 of *kinesinB* (TgGT1_273560) was amplified using primers S12 and AS12, and ligated with pTKO2_II_3_MyoF_mEmeraldFp (Heaslip *et al.,* 2016) via *Eco*RI and *Bgl*II to generate the plasmid pTKO2_C-replace_TgKinesinB_Step1, which lacks a stop codon at the 3’ end of Exon 22. Next, the genomic fragment spanning the 3’ end of *kinesinB* Intron 20, Exon 21 and the 5’ end of Intron 21 was amplified using primers S13 and AS13, and ligated with pTKO2_C-replace_TgKinesinB_Step1 via *Avr*II and *Not*I to generate the plasmid pTKO2_C-replace_TgKinesinB_mEmeraldFP. Lastly, the *mNeonGreenFP* coding sequence was amplified using primers S10 and AS10 with the plasmid pmNeonGreenFP-N1 as a template, and ligated with pTKO2_C-replace_TgKinesinB_mEmeraldFP via *Bgl*II and *Afl*II to generate the 3’ endogenous tagging plasmid pTKO2_C-replace_TgKinesinB_mNeonGreenFP.

pTKO2_II_mCherry_TgKinesinA_5’U_3’UnoHXG, to use as the repair plasmid in conjunction with the *kinesinA* CRISPR/Cas9 plasmid (pU6-Universal_KA_PS, described below) to generate *kinesinA* KO parasites: The ~1.1 kb region immediately upstream of the *kinesinA* locus was amplified using primers S14 and AS14, and ligated with pTKO2_II_mCherryFP (Liu *et al.,* 2013) via *Not*I and *Eco*RV to generate the plasmid pTKO2_II_mCherry_TgKinesinA_5’U. The ~1.5 kb region immediately downstream of the *kinesinA* locus was amplified using primers S15 and AS15 and ligated with pTKO2_II_mCherry_TgKinesinA_5’U via *Nhe*I and *Hpa*I to generate the plasmid pTKO2_II_mCherry_TgKinesinA_5’U_3’U. This plasmid was digested with *Nhe*I and *Spe*I and re-ligated via compatible overhangs to remove the *HXG* cassette to permit selection of *kinesinA* KO parasites with 6-thioxanthine.

pU6-Universal_KA_PS (for Cas9 expression and guide RNA transcription) to use in conjunction with the pTK02_II_mCherry_TgKinesinA_5’U_3’UnoHxG plasmid described above to generate *kinesinA* KO parasites: A protospacer was designed to target the *kinesinA* locus, and checked for possible off-target effects with the *T. gondii* genome sequence (ToxoDB release 28; http://toxodb.org/toxo/ (Gajria *et al.,* 2008; Aurrecoechea *et al.,* 2013)) using ProtoMatch v1.0 (script kindly provided by Sebastian Lourido, Whitehead Institute, Boston, MA). The protospacer was generated by solubilizing primers S16 and AS16 in Duplex Buffer (100 mM potassium acetate, 30 mM HEPES, pH 7.5; filter sterilized) to a final concentration of 200 μM. 20 μL of each of these solubilized oligos was combined and phosphorylated using T4 Polynucleotide Kinase (NEB# M0201) in T4 DNA Ligase Buffer (NEB# B0202) at 37°C for 30 min. The T4 Polynucleotide Kinase was inactivated and protospacer oligos were duplexed by incubating the mixture in a metal microcentrifuge tube block set to 100°C, then allowed to slowly cool to 25°C over 4-16 h. The duplexed protospacer was dialyzed for 20 min using 0.025 μm filter membrane (Millipore Cat# VSWP02500), recovered, and ligated with the plasmid pU6-Universal ((Sidik *et al.,* 2014), Addgene ID# 52694) via *Bsa*I to generate the plasmid pU6-Universal_KA_PS. The protospacer was verified by sequencing with primers S17 and S18.

### Generation of knock-in, endogenously tagged, knock-out, complemented, and transgenic parasites

EGFP-tagged APR1 transgenic parasites: RH parasites (~1 × 10^7^) were electroporated with 30 μg of plasmid coding for ptubA1-APR1-EGFP, and selected with 20 μM chloramphenicol until the population was drug resistant.

*APR1-mCherryFP* knock-in parasites: approximately 1 × 10^7^ *RHΔku80* parasites were electroporated with 50 μg of pTKO2_V-TgAPR1-mCherryFP linearized with *Not*I and selected with 25 μg/mL mycophenolic acid and 50 μg/mL xanthine for three passages, and sorted by FACS for EGFP(-) parasites to reduce the number of parasites in the population that had not undergone double homologous recombination. (The backbone of the pTKO2_V-TgAPR1-mCherryFP plasmid contains an EGFP expression cassette and can therefore help in the identification of single homologous recombinants.) Clones were screened by fluorescence for an mCherryFP(+) ring at the apical end of the parasites, and confirmed with diagnostic genomic PCRs to have the *apr1* endogenous locus replaced by a LoxP-flanked APR1-mCherryFP expression cassette. One clone (C11) was further verified by Southern blotting; this clone was used the subsequent generation of *Δapr1* parasites.

*Δapr1* parasites: *APR1-mCherryFP* knock-in parasites (~1 × 10^7^) were electroporated with 25 μg of pmin-Cre-EGFP (Heaslip *et al.,* 2010), selected with 80 μg/mL of 6-thioxanthine for two passages, and screened for the loss of mCherryFP fluorescence. Clones were confirmed by diagnostic genomic PCRs. One clone (C11:F1) was further verified by Southern blotting and used in all experiments reported here.

*Δapr1:APR1-mCherryFP* complemented parasites: *Δapr1* parasites (~1 × 10^7^) were electroporated with 40 μg of pTKO2_V-TgAPR1-mCherryFP linearized with *Not*I, and selected with 25 μg/mL mycophenolic acid and 50 μg/mL xanthine for two passages. The resultant population expressed cytosolic EGFP in addition to APR1-mCherryFP, indicating that the plasmid had integrated randomly into the genome and not by double homologous recombination at the endogenous *apr1* locus.

3’ endogenously tagged *KinesinA-mNeonGreenFP* parasites: *RHΔku80* parasites (~1 × 10^7^) were electroporated with 50 μg of pTKO2_C-replace_TgKinesinA_mNeonGreenFP linearized with *Kpn*I, and selected with 25 μg/mL mycophenolic acid and 50 μg/mL xanthine for two passages until the parasite population became drug resistant. Clones were tested with diagnostic genomic PCRs to confirm that a single homologous recombination event had occurred at the 3’ end of the *kinesinA* genomic locus to include an mNeonGreenFP expression cassette. One clone (clone 1) of the *KinesinA-mNeonGreenFP* parasites was further verified by Southern blotting and used for generating the *ΔkinesinA* parasites.

*KinesinA-mNeonGreenFP:ptubA1-APR1-mCherryFP* transgenic parasites: *KinesinA-mNeonGreenFP* parasites (~1 × 10^7^) were electroporated with 50 μg of ptubA1-APR1-mCherryFP, and selected with 20 μΜ chloramphenicol for three passages until the parasite population became drug resistant. The population was screened for mCherryFP fluorescence, and one clone (clone G4) was used in the experiments reported here.

*Δapr1:KinesinA-mNeonGreenFP* endogenously tagged parasites: This line was generated as described above for the *KinesinA-mNeonGreenFP* parasites, except that the *Δapr1* parasites were used as the parental strain. One clone (clone 8) was further verified by Southern blotting and used in the experiments reported here.

*ΔkinesinA* parasites: *KinesinA-mNeonGreenFP* parasites (~1 × 10^7^) were electroporated with a mixture of 40 μg of pTKO2_II_mCherry_TgKinesinA_5’U_3’UnoHXG linearized with *NotI* and *Hpa*I (as the repair template) and 40 μg of pU6-Universal_KA_PS (for Cas9 expression and guide RNA transcription). Parasites were selected with 80 μg/mL 6-thioxanthine for four passages until the parasite population became drug resistant. The population was then sorted by FACS to enrich for mNeonGreenFP(-) parasites. Clones were individually screened for the loss of mNeonGreenFP fluorescence and by diagnostic genomic PCRs. One *ΔkinesinA* clone (clone 8) was further verified by Southern blotting and used for generating the *ΔkinesinAΔapr1* parasites. In addition, the *kinesinA* locus was also amplified using S19 and AS17, which anneal outside of the *kinesinA* 5’ and 3’ flanking regions cloned into the *kinesinA* knockout plasmid. The PCR product was gel purified, and sequenced using primers S20, AS18, S21 and AS19 to confirm the loss of the entire *kinesinA* genomic locus in the *ΔkinesinA* parasites.

*ΔkinesinA:APR1-mCherryFP* knock-in parasites: This line was generated as described above for the *APR1-mCherryFP* knock-in parasites, except that the *ΔkinesinA* parasites were used as the parental strain. One clone (clone 1) was further verified by Southern blotting and used in the experiments reported here.

*ΔkinesinAΔapr1* parasites: *ΔkinesinA:APR1-mCherryFP* knock-in parasites (~1 × 10^7^) were electroporated with 25 μg of pmin-Cre-eGFP_Gra-mCherry (Liu *et al.,* 2016), selected with 80 μg/mL of 6-thioxanthine for three passages, and screened for the loss of mCherryFP fluorescence. Clones were confirmed by diagnostic genomic PCRs. One clone (clone 1) was further verified by Southern blotting and used in the experiments reported here.

*ΔkinesinAΔapr1:APR1-mCherryFP* complemented parasites: This line was generated as described above for the *Δapr1:APR1-mCherryFP* complemented parasites, except that the *ΔkinesinAΔapr1* parasites were used as the parental strain.

3’ endogenously tagged *KinesinB-mNeonGreenFP* parasites: *RHΔku80* parasites (~1 × 10^7^) were electroporated with 50 μg of pTKO2_C-replace_TgKinesinB_mNeonGreenFP linearized with *Xho*I, and selected with 25 μg/mL mycophenolic acid and 50 μg/mL xanthine for two passages until the population became drug resistant. Clones, including clone 2 that was used in the experiments reported here, were tested with diagnostic genomic PCRs to confirm that a single homologous recombination event had occurred at the 3’ end of the *kinesinB* genomic locus to include an mNeonGreenFP expression cassette.

Endogenously tagged *RNG1-mCherryFP* parasites: *RHΔku80* and *ΔkinesinAΔapr1* parasites (~1 × 10^7^) were each electroporated with 25 μg of the endogenous tagging plasmid RNG1-LIC-mCherryFP (a kind gift from Dr. Naomi Morrissette, University of California, Irvine, (Tran *et al.,* 2010)) linearized within the 5’ UTR fragment with EcoRV. Parasites were selected with 1 μΜ pyrimethamine for two passages until the population became drug resistant. One clone of each parasite line (clone 1 for *RHΔku80:RNG1-mCherryFP* and clone 5 for *AkinesinAAapr1:RNG1-mCherryFP)* was tested with diagnostic genomic PCRs and Southern blotting to confirm that the *rng1* locus had been fused with the CDS for mCherryFP.

### FACS and parasite cloning

Fluorescence activated cell sorting was performed using an AriaII flow cytometer (BD Biosciences, San Jose, CA) driven by FACSDiva software at the Indiana University Bloomington Flow Cytometry Core Facility (Bloomington, IN). To subclone parasites by limiting dilution, 3-25 parasites with the desired fluorescence profile were sorted per well of a 96-well plate containing confluent HFF monolayers. Wells were screened 7-9 days after sorting for single plaques.

### Southern blotting

Southern blotting was performed as previously described (Liu *et al.,* 2013; Liu *et al.,* 2016) with probes synthesized using components based on the NEBlot Phototope Kit (New England BioLabs, Cat# N7550) and detected using components based on the Phototope-Star detection kit (New England BioLabs, Cat# N7020). All *T. gondii* genomic DNA (gDNA) was prepared from freshly egressed parasites and extracted using the Wizard Genomic DNA Purification kit (Cat# A1120, Promega, Madison, WI).

To probe and detect changes in the *kinesinA* genomic locus in 3’ endogenously tagged *KinesinA-mNeonGreenFP* parasites as well as the RHΔ*ku80*, *ΔkinesinA, ΔkinesinA:APR1-mCherryFP* and *ΔkinesinAΔapr1* parasites, 5 μg of gDNA was digested with *Nco*I. A CDS probe (311 bp) specific for Exon 8 of *kinesinA* was amplified from *T. gondii* RHΔ*ku80* gDNA using primers S22 and AS20, and used as a template in probe synthesis. A probe specific for the region downstream of the *kinesinA* genomic locus (467 bp) was released from plasmid pTKO2_II_mCherry_TgKinesinA_3’U by *Hpa*I and *Sac*I-HF double digestion and used as a template in probe synthesis.

To probe and detect changes in the *apr1* genomic locus in *APR1-mCherryFP* knock-in parasites as well as the RHΔ*ku80*, *Δapr1, Δapr1:KinesinA-mNeonGreenFp* and *ΔkinesinAΔapr1* parasites, 5 μg of gDNA was digested with *Hind*III. A CDS probe (569 bp) specific for *apr1* was amplified from *T. gondii* RHΔ*ku80* gDNA using primers S4 and AS21 and used as a template in probe synthesis. A probe specific for the region upstream of the *apr1* genomic locus (913 bp) was released from plasmid pTKO2_V-TgAPR1-mCherryFP by *Not*I-HF and *Eco*RI-HF double digestion and used as a template in probe synthesis.

To probe and detect changes in the *rng1* (TgGT1_243545; 49.m03355) genomic locus in RH*Δku80* and *ΔkinesinAΔapr1* parasites whose copy of RNG1 was endogenously tagged with mCherryFP, 5 μg of gDNA was digested with *Stu*I and *Mfe*I*-*HF. A probe (454 bp) that anneals upstream of the *rng1* genomic locus was amplified from *T. gondii* RH*Δku80* gDNA using primers S23 and AS22 and used as a template in probe synthesis.

### Plaque assay

Plaque assays were performed as previously described with some modifications (Liu *et al.,* 2016). A total of 50, 100 or 200 freshly egressed parasites were added to each well of a 12- well plate seeded with a confluent HFF monolayer, and incubated undisturbed at 37°C for 7 days. infected monolayers were then fixed with 3.7% (v/v) formaldehyde at 25°C for 10 min, washed with DPBS, stained with 2% (w/v) crystal violet, 20% (v/v) methanol in DPBS for 15 min, gently rinsed with ddH_2_O and air dried.

### Morphometric analysis

Morphometric analysis was performed as previously described with some modifications (Barkhuff *et al.,* 2011). Freshly egressed parasites were centrifuged at 1,000 × g, resuspended in DPBS, allowed to adhere to MatTek dishes coated with Cell-Tak (Cat# 354240, Corning, Oneonta, NY) at 25°C for 30 min, washed gently three times with DPBS and imaged by differential interference (DIC) microscopy. The widest point along the longitudinal and transverse axes of the extracellular parasites was measured using FIJI v.2.0.0-rc-44/1.50e ((Schindelin *et al.,* 2012); http://fiii.se/). Results were calculated from three independent experiments, with 100 parasites per parasite line measured for each experiment.

### Sonieation for generating parasites with radially spread cortical microtubules

To count the number of cortical microtubules that *ΔkinesinAΔapr1* parasites were capable of making, ~1 × 10^7^ freshly egressed, extracellular *ΔkinesinAΔapr1* parasites were centrifuged at 1,000 × *g,* resuspended in 1 mL of DPBS and sonicated for 15 s. The suspension was then centrifuged at 21,000 × *g* for 5 min, with the resulting pellet resuspended in ~10 μL of the supernatant, added to a poly-L-lysine coated coverslip, and then fixed with formaldehyde and processed for immunofluorescence as described below.

### Deoxycholate extraction of extracellular parasites

To label the cortical microtubules in detergent extracted extracellular parasites, ~1 × 10^7^ parasites were centrifuged at 1,000 x *g* and resuspended in ~10 μL of culture medium, spotted onto clean Parafilm and overlaid with a 22 × 22 mm cleaned glass coverslip coated with poly-L-lysine (P8920; SIGMA-Aldrich, St. Louis, MO) for 30-45 min. Parasites were extracted with 10 mM sodium deoxycholate (D6750, SIGMA) in PBS for 15 min and fixed with 3.7% (v/v) formaldehyde in PBS for 15 min and processed for immunofluorescence as described below.

### Immunofluorescence assay

All immunofluorescence labeling steps were performed at room temperature unless otherwise indicated.

For intracellular parasites, *T. gondii-infected* HFF monolayers growing in a 3.5 cm glass-bottom dish were fixed in 3.7% (v/v) formaldehyde in DPBS for 15 min, permeabilized with 0.5% (v/v) Triton X-100 (TX-100) in DPBS for 15 min, blocked in 1% (w/v) BSA in DPBS for 30-60 min, followed by antibody labelling (see below). Dishes were incubated in primary antibodies for 30-60 min followed by incubation in secondary antibodies for 30-60 min unless otherwise noted. Primary antibodies and dilutions used were as follows: rabbit anti-IMC1, 1:1,000 (a kind gift from Dr. Con Beckers, University of North Carolina, Chapel Hill) (Mann and Beckers, 2001); mouse anti-ISP1, 1:1,000 (a kind gift from Dr. Peter Bradley, University of California, Los Angeles) (Beck *et al.,* 2010); mouse anti-MIC2 6D10, 1:1,000 (a kind gift from Dr. Vern Carruthers, University of Michigan, Ann Arbor) (Carruthers *et al.,* 2000), rabbit anti-Tgβ2-tubulin, 1:1,000 and incubation overnight at 8°C (a kind gift from Dr. Naomi Morrissette, University of California, Irvine) (Morrissette and Sibley, 2002b). Secondary antibodies and dilutions used were: goat anti-rabbit IgG Cy3, 1:1,000 (Cat# 111-165-144, Jackson ImmunoResearch); goat anti-mouse IgG Alexa 568, 1:1,000 (Cat# A11031, Molecular Probes); and goat anti-mouse IgG Alexa 488, 1:1,000 (Cat# A11029, Molecular Probes).

For labeling of microtubules in extracellular parasites, after fixation, coverslips were blocked with 1% (w/v) bovine serum albumin (BSA) for 30-60 min, and incubated first with a mixture of mouse anti-α-tubulin and anti-β-tubulin primary antibodies (Cat# T6074 and T5293, respectively; SIGMA) at 1:1,000 for 30-60 min followed by goat anti-mouse IgG Cy3 secondary antibody (Cat# A10521, Molecular Probes) at 1:1,000 for 30-60 min. For samples co-labelled to visualize the parasite cortex, coverslips were additionally incubated with rabbit anti-IMC1 primary antibody (Mann and Beckers, 2001) at 1:1,000 followed by goat anti-rabbit IgG Alexa 488 (Cat# A11008, Life Technologies/Molecular Probes, Eugene, OR) at 1:1,000.

### 2D motility (trail deposition) assay

Trail deposition assays were performed as previously described with minor modifications (Dobrowolski and Sibley, 1996; Carey *et al.,* 2004; Heaslip *et al.,* 2011). 9 × 10^5^ freshly egressed parasites were pelleted at 1,000 x *g,* resuspended in 150 μL of phenol red-free CO_2_- independent medium (Life Technologies-Gibco, SKU#RR060058) and added to 35 mm dishes (#1.5) with a 20 or 14 mm microwell (Cat# P35G-1.5-20-C or Cat# P35G-1.5-14-C, MatTek, Ashland, MA) that had been pre-coated with Cell-Tak (Cat# 354240, Corning, Oneonta, NY). Parasites were allowed to glide at 37°C for 30 min. The samples were fixed with 3.7% (v/v) formaldehyde in DPBS at 25°C for 10 min, washed three times with DPBS and blocked in 1% (w/v) BSA in DPBS for 15 min. Trails were labelled with rabbit anti-P30 (SAG1) antibody (a kind gift from Dr. Lloyd Kasper, Darthmouth College, MA) at 1:100 dilution for 15 min, washed three times with DPBS, incubated in goat anti-mouse IgG Alexa 488 for 15 min and washed three times with DPBS prior to imaging.

### Invasion assays

Invasion assays were performed as previously described (Carey *et al.,* 2004; Mital *et al.,* 2005) with some modifications. Briefly, monolayers were inoculated with equal numbers of freshly egressed parasites, and incubated under conditions that would allow the parasites to invade. The infected monolayers were then fixed and incubated with an antibody that recognizes an epitope of *Toxoplasma* <u>s</u>urface <u>a</u>ntigen 1 (TgSAG1), followed by a secondary antibody that labelled extracellular (non-invaded) parasites red. To distinguish these from parasites that have invaded, the infected host cell monolayers were then permeabilized and incubated with the same anti-TgSAG1 antibody followed by a secondary antibody that labelled both extracellular and intracellular parasites green. Parasites that were labelled red were scored as extracellular, and parasites that were only labelled green were scored as intracellular.

More specifically, 1 × 10^7^ freshly egressed parasites were used to infect a MatTek dish seeded with a ~90% confluent monolayer of BS-C-1 cells. The dishes were incubated at 37°C for 1 hr, washed with DPBS and fixed with 3.7% (v/v) formaldehyde and 0.06% (v/v) glutaraldehyde for 15 min. The samples were then blocked with 1% (w/v) BSA in DPBS for 30-60 min and incubated with mouse anti-TgSAG1 antibody (Cat# 11-132, Argene) at 1:1,000 dilution for 30 min, followed by goat anti-mouse IgG Alexa 568 at 1:1,000 dilution for 30 min. Cells were permeabilized with 0.5% (v/v) TX-100 in DPBS for 15 min, blocked with 1% (w/v) BSA in DPBS for 30-60 min, incubated in mouse anti-TgSAG1 at 1:1,000 dilution for 30 min, followed by goat anti-mouse IgG Alexa 488 at 1:1,000 dilution for 30 min. The samples were imaged at low magnification (Olympus 10X UplanSApo, NA = 0.4), for a total of ten full field images per sample, for each of three independent experiments. Fields were randomly selected using the Alexa 488 channel.

For semi-automated quantification of invaded parasites *(i.e.,* parasites that were labeled with anti-TgSAG1/goat-anti-mouse-Alexa488 but not with anti-TgSAG1/goat-anti-mouse-Alexa568), images were processed using FIJI (ImageJ, v.2.0.0-rc-44/1.50e). The Alexa 488 and Alexa 568 channels were separated, and segmented using the Otsu algorithm for thresholding to create a binarized image followed by watershed separation. The number of intracellular parasites was calculated by subtracting the extracellular parasites (Alexa 568 channel) from the parasites in the Alexa 488 channel, using a size filter of 5-40 μm^2^ and exclusion of objects close to the edges of the image to prevent edge artifacts. Segmentation and counting were manually verified for each image. Pairwise comparisons were made with unpaired, two-tailed Student’s t-tests using GraphPad Prism v. 6.01 (La Jolla, CA).

### Microneme secretion (excretory/secretory antigen assay)

Freshly egressed parasites were harvested and resuspended at a final concentration of 8.0 × 10^8^ parasites/mL in DMEM supplemented with 1% (v/v) CCS. Excreted/secreted antigen preparation was performed by incubating 3.6 × 10^7^ parasites (adjusting the final volume to 90 μL) at 37°C for 7 min in the presence of 1% (v/v) ethanol to assess induced secretion. Tubes were placed on ice for 5 min immediately afterwards, and centrifuged at 1,000 × *g* for 10 min at 4°C. 75 μL of the supernatant was removed and centrifuged again, and 4X sample buffer containing reducing agent was added to 45 μL of the second supernatant. The pellet was washed with DPBS and resuspended in 1X sample buffer containing reducing agent. Samples were boiled for 10 min and resolved using NuPAGE 4-12% Bis-Tris gels, and blotted by wet transfer to nitrocellulose membrane. Membranes were processed for western blotting using the BM Chemiluminescence Western Blotting Kit (Cat# 11520709001, Roche Diagnostics, Indianapolis, IN) according to the manufacturer’s instructions. Briefly, membranes were blocked with 1% (v/v) blocking solution in TBS, probed with mouse mAb anti-TgMIC2 6D10 (1:5,000), in 0.5% (v/v) blocking solution in TBS, washed with TBS-T, and probed with horseradish peroxidase-labelled anti-mouse/rabbit IgG secondary antibody in 0.5% (v/v) blocking solution in TBS. Blots were exposed to chemiluminescence substrate solutions A and B and exposed to film.

### Three-dimensional structured-illumination microscopy (3D-SIM)

3D-SIM image stacks were acquired using an OMX imaging station (GE Healthcare / Applied Precision, Seattle, WA). A 100X oil immersion lens (NA = 1.40) with immersion oil at a refractive index of 1.516 or 1.518 was used; stacks were collected with a z-spacing of 0.125 μm. Images were deconvolved using the point spread functions and software supplied by the manufacturer. Contrast levels were adjusted to better visualize the weak signal without oversaturating the strong signal. All samples were prepared in 35 mm dishes (#1.5) with a 20 or 14 mm microwell (Cat# P35G-1.5-20-C or Cat# P35G-1.5-14-C, MatTek, Ashland, MA), in phenol red-free CO_2_-independent medium (Life Technologies-Gibco, SKU#RR060058) for live imaging and DPBS for fixed samples.

### Wide-field deconvolution microscopy

Image stacks were acquired at 37°C using a DeltaVision imaging station (GE Healthcare / Applied Precision) fitted onto an Olympus IX-70 inverted microscope base. A 60X silicone oil immersion lens (Olympus 60X U-Apo N, NA 1.3) with an auxiliary magnification of 1.5X, or 100X oil immersion lens (Olympus 100X UPLS Apo, NA = 1.40) with immersion oil at a refractive index of 1.524 was used for imaging. 3D image stacks were collected with a z-spacing of 0.3 μm. Images were deconvolved using the point spread functions and software supplied by the manufacturer. Contrast levels were adjusted to better visualize the weak signal without oversaturating the strong signal.

### Electron microscopy

For negative staining of whole mount, detergent-extracted parasites, freshly egressed parasites were harvested and gently washed with 600 μL of phenol red-free CO_2_-independent medium. The parasite pellet was then centrifuged at 3,600 × *g* for 4 min and resuspended in a final volume of 40 μL of the above buffer with or without 20 μΜ A23187, and incubated at 37°C for 5 min. 4 μL of the parasite suspension was then spotted onto hexagonal thin-bar 600-mesh copper grids coated with a carbon film, and incubated at 25°C for 8 min in a humid chamber. The grids were then inverted onto 50 μL of 0.5% (v/v) Triton X-100 in ddH_2_O spotted onto a Teflon sheet, incubated for 3 min, followed by negative staining with 2% (w/v) phosphotungstic acid, pH 7.3 and 0.002% (v/v) Triton X-100 for 3 min. Samples were imaged at a calculated magnification of 20,477X as calibrated using a replica diffraction grating (Ted Pella, Redding, CA) on a JEOL JEM 1010 transmission electron microscope (JEOL, Peabody, MA) operating at 80 keV, with a 100 μm condenser aperture and a 100 μm objective aperture. Images were acquired using a MegaScan 794 CCD camera and DigitalMicrograph v.1.72.53 (Gatan, Pleasanton, CA).

Immunoelectron microscopy on transgenic parasites expressing EGFP-tagged TgAPR1 was performed as described in (Nagayasu *et al.,* 2016).

## Acknowledgements

We thank Drs. Con Beckers (University of North Carolina, Chapel Hill) for the rabbit anti-TgIMC1 antibody, Peter Bradley (University of California, Los Angeles) for the mouse anti-TgISP1 antibody, Vern Carruthers (University of Michigan) for the *RHΔku80Δhx* parasites and mouse mAb 6D10 anti-TgMIC2 antibody, Richard Day (Indiana University School of Medicine, Indianapolis, IN) for the pmNeonGreenFP-N1 and pmAppleFP-C1 plasmids, Sebastian Lourido (Whitehead Institute, Boston, MA) for the pU6-Universal plasmid, for sharing the ProtoMatch v1.0 program and advice on the CRISPR/Cas9 system and Naomi Morrissette (University of California, Irvine) for the rabbit anti-Tgβ2-tubulin antibody and RNG1-mCherry-LIC-DHFR plasmid. We thank Christiane Hassel of the Indiana University Bloomington (IUB) Flow Cytometry Core Facility for assistance with flow cytometry, Dr. James Powers of the IUB Light Microscopy Imaging Center for assistance and support with light microscopy, and Drs. David Morgan and Barry Stein of the IUB Electron Microscopy Center for advice and support with electron microscopy. We would also like to thank Amanda Rollins and Tiffany Fortney for technical support. This study was supported by an American Heart Association Postdoctoral Fellowship (16POST31330004) awarded to J.M.L., funding from the March of Dimes (6-FY15- 198) and the National Institutes of Health/National Institute of Allergy and Infectious Diseases (R01-AI098686) awarded to K.H., and facility funding from the Indiana Clinical and Translational Sciences Institute to K.H., funded in part by Grant UL1 TR001108 from a National Institutes of Health, National Center for Advancing Translational Sciences, Clinical and Translational Sciences Award.

## Supplemental Figures

**Figure S1. A.** Multiple sequence alignment of TgAPR1 and its homologues. TgAPR1, *Toxoplasma gondii* TgGT1_315510; NCLIV_058200, *Neospora caninum* Liverpool conserved hypothetical protein 058200; HHA_315510, *Hammondia hammondi* strain H.H.34 hypothetical protein 315510; Cc_cyc02019, *Cyclospora cayetanensis* hypothetical protein cyc_02019 OEH75465.1; Em_0011080, *Eimeria mitis* hypothetical protein EMH_0011080 (CDJ32003.1, XP_013354568.1); Ea_CDI84105.1, *Eimeria acervulina* conserved hypothetical protein CDI84105.1 (XP_013246853.1). The multiple sequence alignment was generated using Clustal Omega v. 1.2.4 (Goujon *et al.,* 2010; Sievers *et al.,* 2011) and rendered using BoxShade v. 3.21. Black and grey shading indicate identical and similar residues, respectively. **B.** Cartoon showing organization of the KinesinA protein. Numbers indicate amino acid residue position.

**Figure S2.** Generation of endogenously tagged *RNG1-mCherryFP* parasites.

**A.** Schematic for generating endogenously tagged *RNG1-mCherryFP* parasites and Southern blotting strategy. Endogenous tagging of *rng1* with *mCherryFP* [“endogenously tagged (RNG1-mC)”] was performed in *RHΔku80* and *ΔkinesinAΔapr1* parasites (parental) via single crossover homologous recombination. (dup.), *rng1* sequence partially duplicated as a result of the single crossover homologous recombination event. The positions of the restriction sites and probe (red) annealing upstream of the *rng1* genomic locus used in Southern blotting analysis and the corresponding DNA fragment sizes expected are shown.

**B.** Southern blotting analysis of the *rng1* locus in *RHΔku80* (WT), *RHΔku80:RNG1-mCherryFP* (WT:RNG1-mC), *ΔkinesinAΔapr1* (*ΔΔ*) and *ΔkinesinAΔapr1:RNG1-mCherryFP (ΔΔ:RNG1-mC)* parasites generated as described above. The expected parasite genomic DNA fragment sizes after *Stu*I and *Mfe*I digestion are, for the upstream probe, 6214 bp for the parental *(i.e.,* wild-type *rng1* locus), and 3699 bp for the endogenously tagged line.

**Figure S3**. Montage of transmission electron micrographs of negatively stained, whole mount Triton X-100-extracted *ΔkinesinAΔapr1* parasites. Images of parasites with partially or completely detached conoids are indicated with black boxes. Scale bar, 200 nm.

**Figure S4**. Montage of additional, representative transmission electron micrographs of negatively stained, whole mount Triton X-100-extracted *RHΔku80* (WT) parasites. Images of parasites with partially or completely detached conoids are indicated with black boxes. Scale bar, 200 nm.

**Figure S5**. Montage of transmission electron micrographs of negatively stained, whole mount Triton X-100-extracted *APR1-mCherryFP* knock-in *(APR1-mC)* parasites. Scale bar, 200 nm.

**Figure S6**. Montage of transmission electron micrographs of negatively stained, whole mount Triton X-100-extracted *Δapr1* parasites. Images of parasites with partially or completely detached conoids are indicated with black boxes. Scale bar, 200 nm.

**Figure S7**. Montage of transmission electron micrographs of negatively stained, whole mount Triton X-100-extracted *Δapr1:APR1-mCherryFP* complemented parasites (Δ*apr1:APR1-mC*). Images of parasites with partially or completely detached conoids are indicated with black boxes. Scale bar, 200 nm.

**Figure S8**. Map and sequence of the plasmid ptub_EGFP-mCherryFP.

## Supplemental Tables

**Table S1.** List of primers used in this study.

## References

Andenmatten, N., Egarter, S., Jackson, A.J., Jullien, N., Herman, J.P., and Meissner, M. (2012). Conditional genome engineering in *Toxoplasma gondii* uncovers alternative invasion mechanisms. Nature methods, 125-127.

Aurrecoechea, C., Barreto, A., Brestelli, J., Brunk, B.P., Cade, S., Doherty, R., Fischer, S., Gajria, B., Gao, X., Gingle, A., Grant, G., Harb, O.S., Heiges, M., Hu, S., Iodice, J., Kissinger, J.C., Kraemer, E.T., Li, W., Pinney, D.F., Pitts, B., Roos, D.S., Srinivasamoorthy, G., Stoeckert, C.J., Jr., Wang, H., and Warrenfeltz, S. (2013). EuPathDB: the eukaryotic pathogen database. Nucleic acids research 41, D684-691.

Banaszynski, L.A., Chen, L.C., Maynard-Smith, L.A., Ooi, A.G., and Wandless, T.J. (2006). A rapid, reversible, and tunable method to regulate protein function in living cells using synthetic small molecules. Cell 126, 995-1004.

Bannister, L.H., and Mitchell, G.H. (1995). The role of the cytoskeleton in *Plasmodium falciparum* merozoite biology: an electron-microscopic view. Ann Trop Med Parasitol 89, 105-111.

Barkhuff, W.D., Gilk, S.D., Whitmarsh, R., Tilley, L.D., Hunter, C., and Ward, G.E. (2011). Targeted disruption of TgPhIL1 in Toxoplasma gondii results in altered parasite morphology and fitness. PloS one 6, 23977.

Beck, J.R., Rodriguez-Fernandez, I.A., Cruz de Leon, J., Huynh, M.H., Carruthers, V.B., Morrissette, N.S., and Bradley, P.J. (2010). A novel family of *Toxoplasma* IMC proteins displays a hierarchical organization and functions in coordinating parasite division. PLoS pathogens 6,.

Carey, K.L., Westwood, N.J., Mitchison, T.J., and Ward, G.E. (2004). A small-molecule approach to studying invasive mechanisms of *Toxoplasma gondii*. Proc Natl Acad Sci U S A 101, 7433-7438.

Carruthers, V.B., Giddings, O.K., and Sibley, L.D. (1999a). Secretion of micronemal proteins is associated with *Toxoplasma* invasion of host cells. Cell Microbiol 1, 225-235.

Carruthers, V.B., Moreno, S.N., and Sibley, L.D. (1999b). Ethanol and acetaldehyde elevate intracellular [Ca2+] and stimulate microneme discharge in *Toxoplasma gondii*. The Biochemical journal 342 (Pt 2), 379-386.

Carruthers, V.B., Sherman, G.D., and Sibley, L.D. (2000). The *Toxoplasma* adhesive protein MIC2 is proteolytically processed at multiple sites by two parasite-derived proteases. The Journal of biological chemistry 275, 14346-14353.

Carruthers, V.B., and Sibley, L.D. (1997). Sequential protein secretion from three distinct organelles of *Toxoplasma gondii* accompanies invasion of human fibroblasts. European journal of cell biology 73, 114-123.

Carruthers, V.B., and Sibley, L.D. (1999). Mobilization of intracellular calcium stimulates microneme discharge in *Toxoplasma gondii.* Mol Microbiol 31, 421-428.

Dobrowolski, J.M., and Sibley, L.D. (1996). *Toxoplasma* invasion of mammalian cells is powered by the actin cytoskeleton of the parasite. Cell 84, 933-939.

Egarter, S., Andenmatten, N., Jackson, A.J., Whitelaw, J.A., Pall, G., Black, J.A., Ferguson, D.J., Tardieux, I., Mogilner, A., and Meissner, M. (2014). The toxoplasma Acto-MyoA motor complex is important but not essential for gliding motility and host cell invasion. PloS one 9, e91819.

Fox, B.A., Ristuccia, J.G., Gigley, J.P., and Bzik, D.J. (2009). Efficient gene replacements in *Toxoplasma gondii* strains deficient for nonhomologous end joining. Eukaryot Cell 8, 520-529.

Gajria, B., Bahl, A., Brestelli, J., Dommer, J., Fischer, S., Gao, X., Heiges, M., Iodice, J., Kissinger, J.C., Mackey, A.J., Pinney, D.F., Roos, D.S., Stoeckert, C.J., Jr., Wang, H., and Brunk, B.P. (2008). ToxoDB: an integrated Toxoplasma gondii database resource. Nucleic acids research 36, D553-556.

Goujon, M., McWilliam, H., Li, W., Valentin, F., Squizzato, S., Paern, J., and Lopez, R. (2010). A new bioinformatics analysis tools framework at EMBL-EBI. Nucleic acids research 38, W695-699.

Hakansson, S., Morisaki, H., Heuser, J., and Sibley, L.D. (1999). Time-lapse video microscopy of gliding motility in *Toxoplasma gondii* reveals a novel, biphasic mechanism of cell locomotion. Molecular biology of the cell 10, 3539-3547.

Heaslip, A.T., Dzierszinski, F., Stein, B., and Hu, K. (2010). TgMORN1 is a key organizer for the basal complex of *Toxoplasma gondii.* PLoS pathogens 6, e1000754.

Heaslip, A.T., Nelson, S.R., and Warshaw, D.M. (2016). Dense granule trafficking in Toxoplasma gondii requires a unique class 27 myosin and actin filaments. Molecular biology of the cell 27, 2080-2089.

Heaslip, A.T., Nishi, M., Stein, B., and Hu, K. (2011). The Motility of a Human Parasite, *Toxoplasma gondii,* Is Regulated by a Novel Lysine Methyltransferase. PLoS pathogens 7, e1002201.

Heidemann, S.R., and McIntosh, J.R. (1980). Visualization of the structural polarity of microtubules. Nature 286, 517-519.

Hu, K. (2008). Organizational changes of the daughter basal complex during the parasite replication of *Toxoplasma gondii*. PLoS pathogens 4, e10.

Hu, K., Johnson, J., Florens, L., Fraunholz, M., Suravajjala, S., Dilullo, C., Yates, J., Roos, D.S., and Murray, J.M. (2006). Cytoskeletal Components of an Invasion Machine-The Apical Complex of *Toxoplasma gondii.* PLoS pathogens 2, e13.

Hu, K., Roos, D.S., and Murray, J.M. (2002). A novel polymer of tubulin forms the conoid of *Toxoplasma gondii.* The Journal of cell biology 156, 1039-1050.

Hu, K.S.S.; DiLullo, C.; Roos, D.S.; Murray, J. M.. (2003). Identification of new tubulin isoforms in *Toxoplasma gondii.* ASCB Annual Meeting *Abstract #L460.*

Huynh, M.H., and Carruthers, V.B. (2006). *Toxoplasma* MIC2 is a major determinant of invasion and virulence. PLoS pathogens 2, e84.

Huynh, M.H., and Carruthers, V.B. (2009). Tagging of endogenous genes in a *Toxoplasma gondii* strain lacking Ku80. Eukaryot Cell 8, 530-539.

Huynh, M.H., Rabenau, K.E., Harper, J.M., Beatty, W.L., Sibley, L.D., and Carruthers, V.B. (2003). Rapid invasion of host cells by *Toxoplasma* requires secretion of the MIC2-M2AP adhesive protein complex. EMBO J 22, 2082-2090.

Katris, N.J., van Dooren, G.G., McMillan, P.J., Hanssen, E., Tilley, L., and Waller, R.F. (2014). The apical complex provides a regulated gateway for secretion of invasion factors in Toxoplasma. PLoS pathogens 10, e1004074.

Kremers, G.J., Hazelwood, K.L., Murphy, C.S., Davidson, M.W., and Piston, D.W. (2009). Photoconversion in orange and red fluorescent proteins. Nature methods 6, 355-358.

Leung, J.M., Rould, M.A., Konradt, C., Hunter, C.A., and Ward, G.E. (2014). Disruption of TgPHIL1 Alters Specific Parameters of Toxoplasma gondii Motility Measured in a Quantitative, Three-Dimensional Live Motility Assay. PloS one 9, e85763.

Liu, J., He, Y., Benmerzouga, I., Sullivan, W.J., Jr., Morrissette, N.S., Murray, J.M., and Hu, K. (2016). An ensemble of specifically targeted proteins stabilizes cortical microtubules in the human parasite Toxoplasma gondii. Molecular biology of the cell 27, 549-571.

Liu, J., Wetzel, L., Zhang, Y., Nagayasu, E., Ems-McClung, S., Florens, L., and Hu, K. (2013). Novel thioredoxin-like proteins are components of a protein complex coating the cortical microtubules of Toxoplasma gondii. Eukaryot Cell.

Lourido, S., and Moreno, S.N. (2015). The calcium signaling toolkit of the Apicomplexan parasites Toxoplasma gondii and Plasmodium spp. Cell Calcium 57, 186-193.

Lovett, J.L., Marchesini, N., Moreno, S.N., and Sibley, L.D. (2002). *Toxoplasma gondii* microneme secretion involves intracellular Ca(2+) release from inositol 1,4,5-triphosphate (IP(3))/ryanodine-sensitive stores. The Journal of biological chemistry 277, 25870-25876.

Mann, T., and Beckers, C. (2001). Characterization of the subpellicular network, a filamentous membrane skeletal component in the parasite *Toxoplasma gondii*. Molecular and biochemical parasitology 115, 257-268.

Mital, J., Meissner, M., Soldati, D., and Ward, G.E. (2005). Conditional expression of *Toxoplasma gondii* apical membrane antigen-1 (TgAMA1) demonstrates that TgAMA1 plays a critical role in host cell invasion. Molecular biology of the cell 16, 4341-4349.

Morrissette, N.S., Mitra, A., Sept, D., and Sibley, L.D. (2004). Dinitroanilines bind alpha-tubulin to disrupt microtubules. Molecular biology of the cell 15, 1960-1968.

Morrissette, N.S., Murray, J.M., and Roos, D.S. (1997). Subpellicular microtubules associate with an intramembranous particle lattice in the protozoan parasite *Toxoplasma gondii.* Journal of cell science 110, 35-42.

Morrissette, N.S., and Sibley, L.D. (2002a). Cytoskeleton of apicomplexan parasites. Microbiol Mol Biol Rev 66, 21-38; table of contents.

Morrissette, N.S., and Sibley, L.D. (2002b). Disruption of microtubules uncouples budding and nuclear division in *Toxoplasma gondii.* Journal of cell science 115, 1017-1025.

Nagayasu, E., Hwang, Y.-c., Liu, J., Murray, J., and Hu, K. (2016). Loss of a doublecortin (DCX) domain containing protein causes structural defects in a tubulin-based organelle of <em>Toxoplasma gondii</em> and impairs host cell invasion. bioRxiv

Nichols, B.A., and Chiappino, M.L. (1987). Cytoskeleton of *Toxoplasma gondii*. J Protozool 34, 217-226.

Nichols, B.A., Chiappino, M.L., and O’Connor, G.R. (1983). Secretion from the rhoptries of *Toxoplasma gondii* during host-cell invasion. J Ultrastruct Res 83, 85-98.

Opitz, C., and Soldati, D. (2002). ‘The glideosome’: a dynamic complex powering gliding motion and host cell invasion by *Toxoplasma gondii.* Mol Microbiol 45, 597-604.

Pidoux, A.L., LeDizet, M., and Cande, W.Z. (1996). Fission yeast pkl1 is a kinesin-related protein involved in mitotic spindle function. Molecular biology of the cell 7, 1639-1655.

Porchet, E., and Torpier, G. (1977). [Freeze fracture study of *Toxoplasma* and *Sarcocystis* infective stages (author’s transl)]. Z Parasitenkd 54, 101-124.

Roos, D.S., Donald, R.G., Morrissette, N.S., and Moulton, A.L. (1994). Molecular tools for genetic dissection of the protozoan parasite *Toxoplasma gondii.* In: Methods in Cell Biology, vol. 45, 27-63.

Russell, D.G., and Burns, R.G. (1984). The polar ring of coccidian sporozoites: a unique microtubule-organizing centre. Journal of cell science 65, 193-207.

Schindelin, J., Arganda-Carreras, I., Frise, E., Kaynig, V., Longair, M., Pietzsch, T., Preibisch, S., Rueden, C., Saalfeld, S., Schmid, B., Tinevez, J.Y., White, D.J., Hartenstein, V., Eliceiri, K., Tomancak, P., and Cardona, A. (2012). Fiji: an open-source platform for biological-image analysis. Nature methods 9, 676-682.

Shaner, N.C., Campbell, R.E., Steinbach, P.A., Giepmans, B.N., Palmer, A.E., and Tsien, R.Y. (2004). Improved monomeric red, orange and yellow fluorescent proteins derived from Discosoma sp. red fluorescent protein. Nat Biotechnol 22, 1567-1572.

Shaner, N.C., Lambert, G.G., Chammas, A., Ni, Y., Cranfill, P.J., Baird, M.A., Sell, B.R., Allen, J.R., Day, R.N., Israelsson, M., Davidson, M.W., and Wang, J. (2013). A bright monomeric green fluorescent protein derived from Branchiostoma lanceolatum. Nature methods 10, 407-409.

Sheffield, H.G., and Melton, M.L. (1968). The fine structure and reproduction of *Toxoplasma gondii.* The Journal of parasitology 54, 209-226.

Shen, B., Brown, K.M., Lee, T.D., and Sibley, L.D. (2014). Efficient gene disruption in diverse strains of Toxoplasma gondii using CRISPR/CAS9. mBio 5, e01114-01114.

Sibley, L.D. (2010). How apicomplexan parasites move in and out of cells. Curr Opin Biotechnol, 592-598.

Sidik, S.M., Hackett, C.G., Tran, F., Westwood, N.J., and Lourido, S. (2014). Efficient genome engineering of Toxoplasma gondii using CRISPR/Cas9. PloS one 9, e100450.

Sievers, F., Wilm, A., Dineen, D., Gibson, T.J., Karplus, K., Li, W., Lopez, R., McWilliam, H., Remmert, M., Soding, J., Thompson, J.D., and Higgins, D.G. (2011). Fast, scalable generation of high-quality protein multiple sequence alignments using Clustal Omega. Mol Syst Biol 7, 539

Suvorova, E.S., Francia, M., Striepen, B., and White, M.W. (2015). A novel bipartite centrosome coordinates the apicomplexan cell cycle. PLoS Biol 13, e1002093.

Tran, J.Q., de Leon, J.C., Li, C., Huynh, M.H., Beatty, W., and Morrissette, N.S. (2010). RNG1 is a late marker of the apical polar ring in *Toxoplasma gondii.* Cytoskeleton (Hoboken) 67, 586-598.

Tran, J.Q., Li, C., Chyan, A., Chung, L., and Morrissette, N.S. (2012). SPM1 stabilizes subpellicular microtubules in *Toxoplasma gondii.* Eukaryot Cell 11, 206-216.

Wu, Y., Chandris, P., Winter, P.W., Kim, E.Y., Jaumouille, V., Kumar, A., Guo, M., Leung, J.M., Smith, C., Rey-Suarez, I., Liu, H., Waterman, C.M., Ramamurthi, K.S., La Riviere, P.J., and Shroff, H. (2016). Simultaneous multiview capture and fusion improves spatial resolution in wide-field and light-sheet microscopy. Optica 3, 897-910.

Yukawa, M., Ikebe, C., and Toda, T. (2015). The Msd1-Wdr8-Pkl1 complex anchors microtubule minus ends to fission yeast spindle pole bodies. The Journal of cell biology 209, 549-562.

